# Altered cortico-subcortical network after adolescent alcohol exposure mediates behavioral deficits in flexible decision-making

**DOI:** 10.1101/2021.03.12.435040

**Authors:** Alexander Gómez-A, Carol A. Dannenhoffer, Amanda Elton, SungHo Lee, Woomi Ban, Yen-Yu Ian Shih, Charlotte A. Boettiger, Donita L. Robinson

**Affiliations:** Bowles Center for Alcohol Studies, University of North Carolina, Chapel Hill, NC, USA; Department of Psychology and Neuroscience, University of North Carolina, Chapel Hill, NC, USA; Biomedical Research Imaging Center, University of North Carolina, Chapel Hill, NC USA; Department of Neurology, University of North Carolina, Chapel Hill, NC, USA; Neuroscience Curriculum, University of North Carolina, Chapel Hill, NC, USA; Department of Psychiatry, University of North Carolina, Chapel Hill, NC, USA

**Author notes:** Corresponding author, Author information: Dr. Donita L. Robinson, Bowles Center for Alcohol Studies, CB 7178, University of North Carolina, NC 27599-7178, USA, 1+.

## Abstract

Behavioral flexibility, the ability to modify behavior according to changing conditions, is essential to optimize decision-making. Deficits in behavioral flexibility that persist into adulthood are one consequence of adolescent alcohol exposure, and another is decreased functional connectivity in brain structures involved in decision-making; however, a link between these two outcomes has not been established. We assessed effects of adolescent alcohol and sex on both Pavlovian and instrumental behaviors and functional connectivity in adult animals to determine associations between behavioral flexibility and resting-state functional connectivity. Alcohol exposure impaired attentional set reversals and decreased functional connectivity among cortical and subcortical regions-of-interest that underlie flexible behavior. Moreover, mediation analyses indicated that adolescent alcohol-induced reductions in functional connectivity within a subnetwork of affected brain regions mediated errors committed during reversal learning. These results provide a novel link between persistent reductions in brain functional connectivity and deficits in behavioral flexibility resulting from adolescent alcohol exposure.

## Introduction

You are heading to dinner at a place near home. Halfway there, it starts raining, and you duck under a shelter. You consider whether you should go back home for an umbrella, wait a few minutes for the rain to stop, or just keep walking to dinner. What do you do? A crucial element of decision-making is flexibility, which implies the ability to adjust behavior in response to both environmental demands and personal factors^1,2^. However, this process requires complementary psychological functions like inhibitory control and working memory, among other components of executive function^1^.

Executive functions recruit subcortical and frontal, parietal and temporal cortical areas^3^, all of which are still developing during adolescence^4^. Nevertheless, there has been a dominant interest in the role of the prefrontal cortex in executive function because of its involvement in “top-down control” (for review see^3^). The prefrontal cortex matures later in development than other cortical regions^2,4^, and this process might be altered by recreational drugs^5^, including alcohol (see^6,7^ for review). Since executive function depends on adequate brain development, and flexible decision-making requires proper executive function, any potential hazard to brain development is detrimental to these processes.

Interest in the consequences of underage binge alcohol drinking has increased recently, accompanied by many studies in animal models demonstrating the harmful and persistent effects that adolescent alcohol exposure can have on brain development and, consequently, on behavior. For example, adolescent intermittent ethanol (AIE) exposure can promote adult alcohol intake in male and female rodents^8,9^, increase anxiety-like behaviors^10^, perpetuate adolescent-typical behavior in adulthood^11^, and impair cognitive processing, especially in tasks that require inhibitory control, working memory and flexible decision-making^6,12,13^. Moreover, AIE exposure increases conditioned approach to reward-associated cues^14,15^ and impairs flexible responses when experimental conditions change^9,16–18^.

Consistent with those findings, we recently reported that AIE exposure shifted conditioned approach away from goal-tracking and toward sign-tracking^19^ during Pavlovian conditioned approach. While the goal-tracking response generally adapts to changing conditions, such as reward omission, sign-tracking behavior is less flexible^20,21^. In additional studies, AIE exposure decreased reversal learning of an attentional set^22^, indicating impaired inhibition of a previously learned response to a reward-predictive cue when the stimulus-response contingency changed. These behavioral tasks are governed by fronto-striatal circuits involved in action selection, and a separate study demonstrated that AIE exposure decreased functional connectivity of these frontostriatal circuits in rats^23^. Together, these data suggest a potential relationship between inflexible sign-tracking and inflexible attentional set strategy after AIE exposure, potentially mediated by changes in functional connectivity among prefrontal and striatal brain circuits involved in action selection. However, the fact that those results came from separate studies impedes the integration of these AIE effects. Here, we addressed this gap by testing the effects of AIE exposure on sensitivity to conditioned reinforcers (Pavlovian conditioned approach) and behavioral flexibility (attentional set-shifting) within the same animals and integrated them with resting-state functional connectivity MRI among frontal cortical and subcortical regions that regulate action selection. Then, using a dimensionality reduction approach, we identified a principal component of functional connections (functional subnetwork) to test potential mediational effects of functional connectivity on behavior.

Based on our previous studies, we hypothesized that AIE exposure promotes greater sensitivity to reward conditioning and inflexible action-selection in adulthood by altering shared underlying neural circuits. Specifically, we predicted that AIE exposure would increase bias to reward-associated cues and impair behavioral flexibility in adulthood when action outcomes change, and that those effects would be mediated by decreased functional connectivity after AIE exposure.

## Results

### Pavlovian Conditioned Approach

Rats were exposed to binge levels of ethanol (5 g/kg/day ethanol intragastrically, 2-days-on/2-days-off, P25-54) or equivalent volumes of water during adolescence, and then allowed to age into adulthood (Fig. 1A). Animals trained on Pavlovian conditioned approach for 20 daily sessions across four weeks, in which a 30-second stimulus (light and lever extension) predicted a sucrose reward. For data analysis, we averaged metrics of conditioned approach across the five last sessions to assess effects of sex or alcohol exposure. Behaviors were categorized as either sign-tracking (lever-directed) or goal-tracking (receptacle-directed; Fig. 1B; Methods). Lever-directed behaviors included the number of lever presses, latency to press the lever within trials, and probability of pressing the lever within sessions. Receptacle-directed behaviors included the receptacle entry elevation score (calculated as the number of entries during a cue presentation minus entries during the 30 seconds before the cue presentation^19,24^), latency to enter the receptacle, and probability of entering the receptacle. Using a generalized linear model and calculating odds ratios (OR) and their corresponding 95% confidence intervals (CI; Methods), we found significant main effects of sex across all sign-tracking behaviors, with females pressing the lever more times, sooner, and with higher probability of any lever press (Fig. 1C; Supplemental figure 1A; full statistical outcomes in Table 1). We also observed that AIE-exposed rats presented a trending decrease in lever press probability (P=0.09). No interaction between exposure and sex was observed for any lever-directed metric (all P>0.05; Table 1). Regarding goal-tracking behaviors, only female rats presented a trend effect on receptacle latencies (P=0.09), showing a slight decrease in female latencies compared to male latencies. No other significant effects of AIE exposure, sex or exposure-by-sex interaction were found for receptacle-directed behavioral metrics (all P>0.05; Fig. 1C; Supplemental Fig. 1B; Table 1). In summary, our data found that females were biased to interact with the cue, independent of adolescent alcohol exposure.

**Figure 1.**
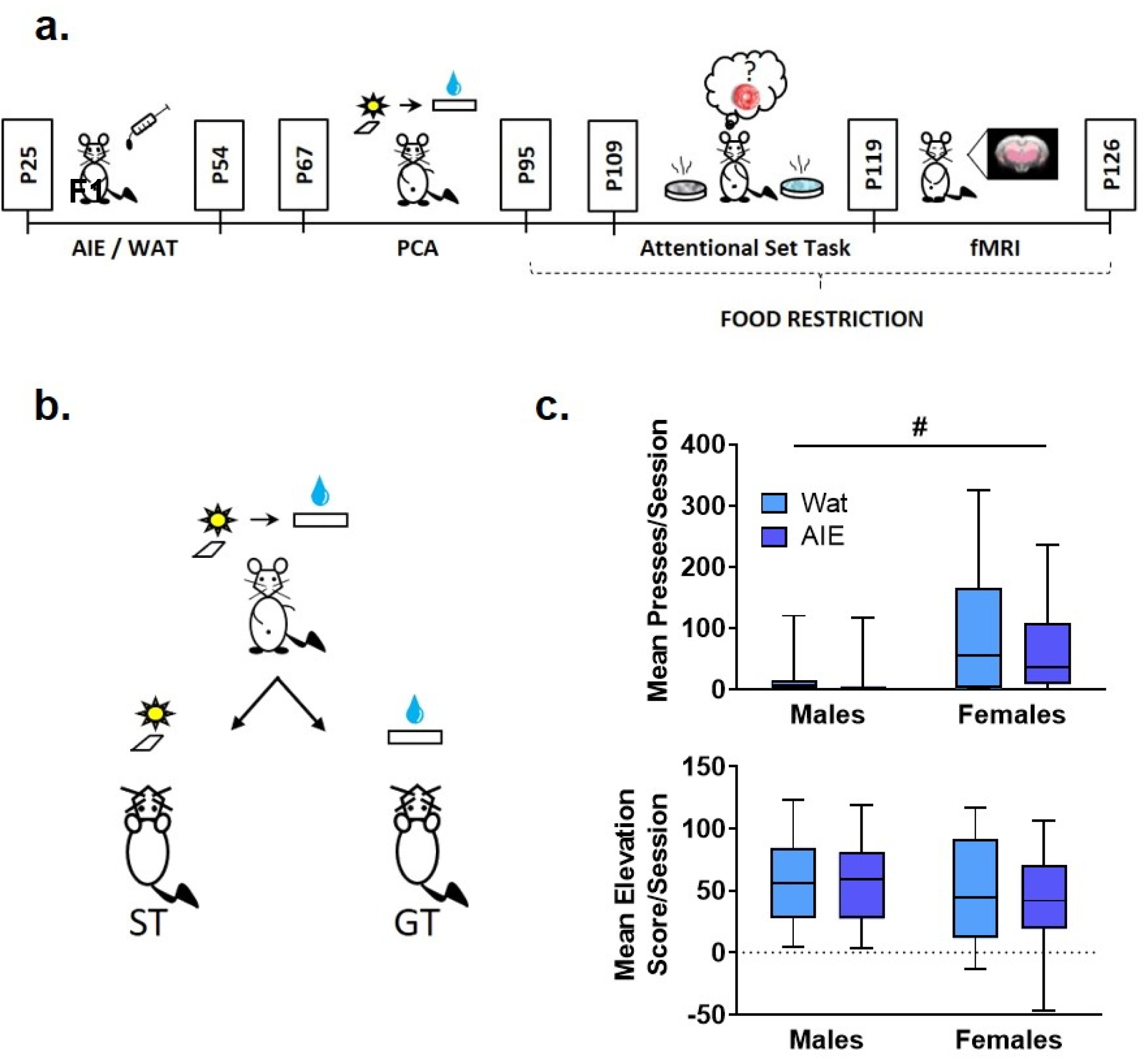
AIE exposure did not affect Pavlovian conditioned approach behavior. **A.** Scheme illustrating the experimental outline used in this study. Male and female rats were exposed to alcohol (5 g/kg) or water during adolescence. On ~P67 rats started behavioral training in Pavlovian conditioned approach (four weeks) followed by training in an attentional set-shifting task (nine days); the attentional set-shift was preceded by two weeks of food restriction. After behavioral training, animals underwent a single session of resting-state fMRI. **B.** Scheme representing phenotypes of interest in Pavlovian conditioned approach. During training, a compound-cue (light/lever) was presented during a 30-second period, followed by 100 μl of 20% sucrose that served as a reward. Sign-tracking (ST) rats preferentially interacted with the cue (light/lever) while goal-tracking (GT) animals preferentially interacted with the reward receptacle. **C. Top:** Sex, but not AIE exposure, promoted ST behavior, reflected in more conditioned lever presses by female rats than males. **Bottom:** No differences were observed in elevation score, a measure of GT behavior. Since the data were not normally distributed, data are presented in box plots showing median (horizontal line), interquartile range (box), and minimum and maximum data values (lower and upper whiskers). # main effect of sex. Detailed statistical analyses are provided in Table 1, additional data from Pavlovian conditioned approach are shown in Supplemental Fig. 1.

**Table 1.** Statistical analysis results of the Pavlovian conditioned approach and the attentional set-shifting task (N=79 rats). AIE exposure and sex effects on conditioned approach (except elevation score) and the attentional set-shifting task metrics were analyzed using a generalized linear model with a Poisson distribution and a log link function. The elevation score (@) analysis was carried out using a two-way ANOVA. * Significant at p ≤ 0.05.

| **I. Pavlovian Conditioning Approach** |  |  |  |  |  |
| --- | --- | --- | --- | --- | --- |
| **Variable** | **Effect** | **P** | **Exp(B)** | **Coefficient Interval** |  |
|  |  |  |  | **Lower** | **Upper** |
| Lever presses | Exposure | 0.39 | 0.508 | 0.105 | 2.453 |
|  | Sex | **<0.01\*** | 4.541 | 1.697 | 12.148 |
|  | Exp. x Sex | 0.70 | 1.387 | 0.251 | 7.66 |
| Lever latency | Exposure | 0.27 | 1.110 | 0.920 | 1.338 |
|  | Sex | **<0.01\*** | 0.721 | 0.590 | 0.883 |
|  | Exp. x Sex | 0.75 | 0.954 | 0.714 | 1.275 |
| Lever probability | Exposure | 0.09 | 0.509 | 0.228 | 1.136 |
|  | Sex | **<0.01\*** | 2.274 | 1.322 | 3.912 |
|  | Exp. x Sex | 0.23 | 1.743 | 0.695 | 4.367 |
| Receptacle entries | Exposure | 0.67 | 0.931 | 0.664 | 1.305 |
|  | Sex | 0.59 | 1.088 | 0.795 | 1.490 |
|  | Exp. x Sex | 0.81 | 0.944 | 0.590 | 1.512 |
| Receptacle latency | Exposure | 0.96 | 0.993 | 0.736 | 1.339 |
|  | Sex | 0.09 | 0.769 | 0.565 | 1.045 |
|  | Exp. x Sex | 0.48 | 1.171 | 0.753 | 1.82 |
| Receptacle probability | Exposure | 0.62 | 0.978 | 0.895 | 1.068 |
|  | Sex | 0.14 | 1.064 | 0.979 | 1.157 |
|  | Exp. x Sex | 0.73 | 0.979 | 0.866 | 1.108 |
| **Variable** | **Effect** | **P** | **F** | **Df** |  |
| Elevation Score<sup>@</sup> | Exposure | 0.41 | 0.658 | 1 |  |
|  | Sex | 0.19 | 1.673 | 1 |  |
|  | Exp. x Sex | 0.57 | 0.321 | 1 |  |
| **II. Attentional Set-shift task** |  |  |  |  |  |
| **a. Acquisition** |  |  |  |  |  |
| **Variable** | **Effect** | **P** | **Exp(B)** | **Coefficient Interval** |  |
|  |  |  |  | **Lower** | **Upper** |
| Total trials acquisition | Exposure | 0.92 | 0.984 | 0.725 | 1.336 |
|  | Sex | 0.26 | 1.174 | 0.885 | 1.558 |
|  | Exp. x Sex | 0.58 | 0.890 | 0.584 | 1.356 |
| Active errors acquisition | Exposure | 0.26 | 0.622 | 0.270 | 1.434 |
|  | Sex | 0.43 | 1.302 | 0.671 | 2.527 |
|  | Exp. x Sex | 0.83 | 1.123 | 0.376 | 3.35 |
| **b. Reversal 1** |  |  |  |  |  |
| Total trials reacquisition | Exposure | 0.65 | 0.951 | 0.761 | 1.188 |
|  | Sex | 0.30 | 0.893 | 0.718 | 1.110 |
|  | Exp. x Sex | 0.26 | 1.198 | 0.875 | 1.642 |
| Active errors reacquisition | Exposure | 0.33 | 0.652 | 0.275 | 1.545 |
|  | Sex | 0.11 | 0.476 | 0.192 | 1.184 |
|  | Exp. x Sex | **0.05\*** | 3.581 | 1.021 | 12.557 |
| Total trials acquisition (R1) | Exposure | 0.66 | 1.066 | 0.799 | 1.422 |
|  | Sex | 0.57 | 0.921 | 0.690 | 1.228 |
|  | Exp. x Sex | 0.55 | 1.132 | 0.753 | 1.701 |
| Active errors acquisition (R1) | Exposure | 0.25 | 1.173 | 0.891 | 1.545 |
|  | Sex | 0.42 | 0.890 | 0.670 | 1.183 |
|  | Exp. x Sex | 0.14 | 1.330 | 0.904 | 1.956 |
| Prepotent errors<br>(R1) | Exposure | 0.08 | 1.525 | 0.950 | 2.447 |
|  | Sex | 0.80 | 1.066 | 0.649 | 1.749 |
|  | Exp. x Sex | 0.39 | 1.320 | 0.699 | 2.496 |
| Regressive errors<br>(R1) | Exposure | 0.95 | 1.021 | 0.532 | 1.960 |
|  | Sex | 0.54 | 0.814 | 0.418 | 1.584 |
|  | Exp. x Sex | 0.60 | 1.277 | 0.5 | 3.258 |
| Initial regressive<br>errors<br>(R1) | Exposure | 0.86 | 1.053 | 0.591 | 1.876 |
|  | Sex | 0.46 | 0.785 | 0.431 | 1.431 |
|  | Exp. x Sex | 0.59 | 1.253 | 0.541 | 2.903 |
| Subsequent<br>regressive errors<br>(R1) | Exposure | 0.93 | 0.961 | 0.376 | 2.458 |
|  | Sex | 0.76 | 0.870 | 0.344 | 2.199 |
|  | Exp. x Sex | 0.67 | 1.329 | 0.355 | 4.977 |
| **c. Reversal 2** |  |  |  |  |  |
| Total trials<br>reacquisition | Exposure | 0.08 | 1.197 | 0.977 | 1.467 |
|  | Sex | 0.98 | 0.997 | 0.813 | 1.224 |
|  | Exp. x Sex | 0.89 | 1.080 | 0.813 | 1.433 |
| Active errors<br>reacquisition | Exposure | **0.03\*** | 1.905 | 1.061 | 3.421 |
|  | Sex | 0.34 | 1.342 | 0.73 | 2.466 |
|  | Exp. x Sex | 0.94 | 0.973 | 0.451 | 2.098 |
| Total trials<br>acquisition<br>(R2) | Exposure | 0.10 | 1.060 | 0.757 | 1.485 |
|  | Sex | **0.04\*** | 1.110 | 0.805 | 1.532 |
|  | Exp. x Sex | 0.26 | 1.298 | 0.828 | 2.034 |
| Active errors<br>acquisition<br>(R2) | Exposure | **<0.01\*** | 1.563 | 1.144 | 2.135 |
|  | Sex | 0.10 | 1.295 | 0.946 | 1.774 |
|  | Exp. x Sex | 0.46 | 1.165 | 0.775 | 1.750 |
| Prepotent errors<br>(R2) | Exposure | **0.02\*** | 1.842 | 1.116 | 3.042 |
|  | Sex | 0.06 | 0.530 | 0.274 | 1.025 |
|  | Exp. x Sex | 0.76 | 1.137 | 0.497 | 2.604 |
| Regressive errors<br>(R2) | Exposure | 0.10 | 1.404 | 0.941 | 2.094 |
|  | Sex | **<0.01\*** | 1.753 | 1.208 | 2.545 |
|  | Exp. x Sex | 0.37 | 1.247 | 0.764 | 2.037 |
| Initial regressive<br>errors<br>(R2) | Exposure | 0.60 | 1.18 | 0.633 | 2.202 |
|  | Sex | 0.22 | 1.433 | 0.802 | 2.558 |
|  | Exp. x Sex | 0.36 | 1.434 | 0.655 | 3.139 |
| Subsequent<br>regressive errors<br>(R2) | Exposure | **0.05\*** | 2.222 | 1.005 | 4.912 |
|  | Sex | **<0.01\*** | 2.828 | 1.335 | 5.994 |
|  | Exp. x Sex | 0.75 | 0.864 | 0.343 | 2.177 |
| **d. Set shift** |  |  |  |  |  |
| Total trials<br>reacquisition | Exposure | 0.35 | 0.871 | 0.649 | 1.168 |
|  | Sex | 0.50 | 0.909 | 0.687 | 1.202 |
|  | Exp. x Sex | 0.56 | 1.131 | 0.745 | 1.716 |
| Active errors<br>reacquisition | Exposure | 0.44 | 0.741 | 0.343 | 1.6 |
|  | Sex | 0.97 | 1.010 | 0.511 | 1.998 |
|  | Exp. x Sex | 0.66 | 1.265 | 0.443 | 3.613 |
| Total trials<br>acquisition<br>(SS) | Exposure | 0.88 | 1.017 | 0.804 | 1.287 |
|  | Sex | 0.97 | 0.997 | 0.793 | 1.251 |
|  | Exp. x Sex | 0.30 | 1.186 | 0.857 | 1.64 |
| Active errors<br>acquisition<br>(SS) | Exposure | 0.42 | 1.203 | 0.762 | 1.899 |
|  | Sex | 0.77 | 0.935 | 0.586 | 1.493 |
|  | Exp. x Sex | 0.16 | 1.552 | 0.833 | 2.891 |

### Attentional set-shifting task

Next, we trained rats on an attentional set-shifting task^25^ (Fig. 1A and 2A) to discriminate between two different odors, one of which was associated with a small piece of food that served as a reward (the behavior paradigm is detailed in Supplemental Table 1 and Methods; adapted from Birrell and Brown^25^). Cups containing rewards were covered with digging media, and animals were first trained to dig to find the reward, then to use odor as a discriminative stimulus to predict the reward independent of the digging medium. Media and odor stimuli are listed in Supplemental Table 1. After acquisition of the discrimination, two reversals were introduced, keeping odor as the relevant dimension (intradimensional reversal), followed by a set-shift phase, where the appropriate discriminant was the digging medium instead of odor (extradimensional shift). We measured the number of trials that animals required to reach the criterion of 6 consecutive correct choices, as well as the total number of errors made. Here, we report results from this task according to each step: Acquisition, Reversal 1, Reversal 2, and Extradimensional Set-Shift (full statistics in Table 1).

**Figure 2.**
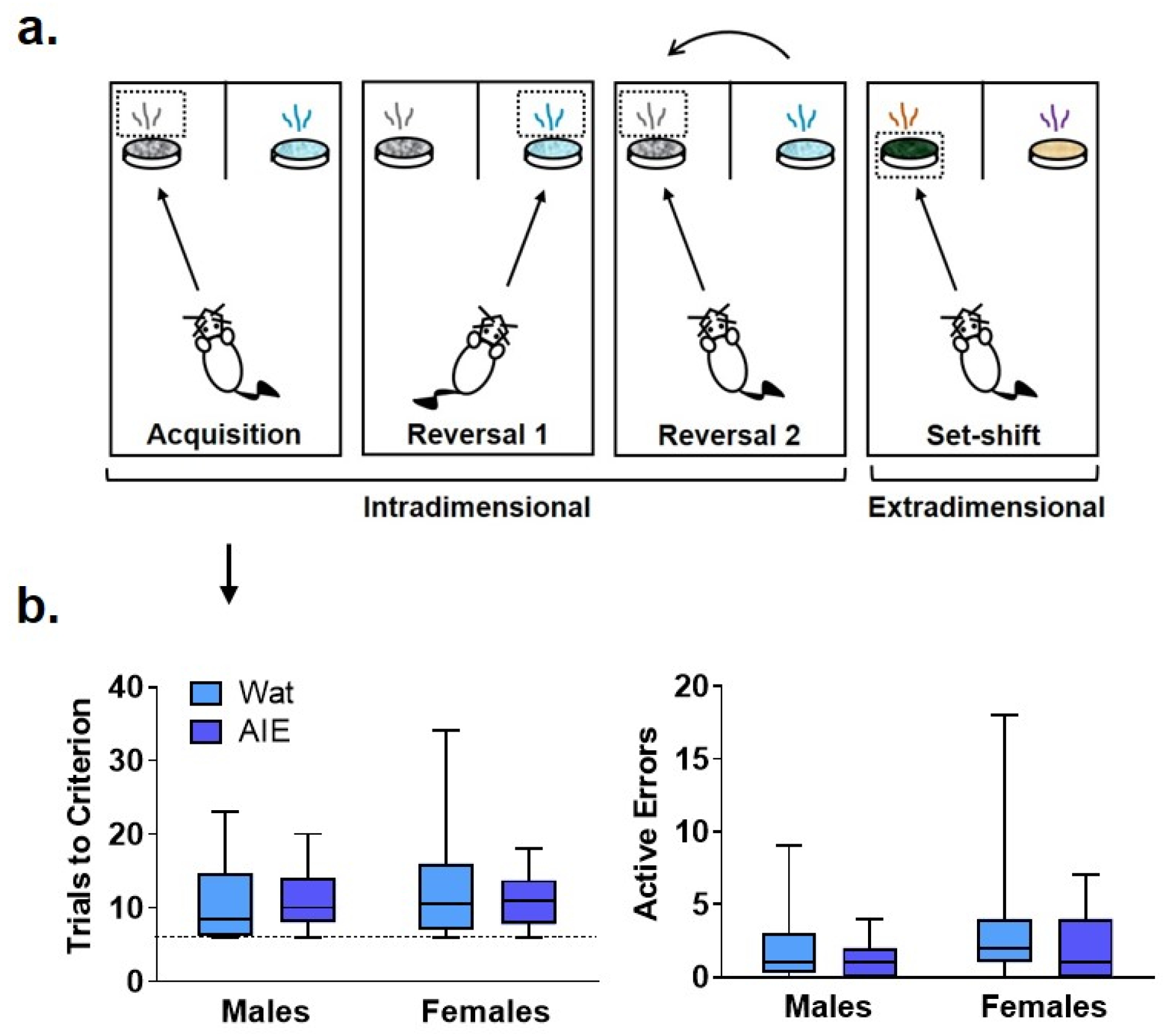
AIE exposure did not impair initial discriminative learning in an attentional set-shifting task. **A.** Scheme of the attentional set-shifting task. We trained rats to discriminate between two different odors, one of which was associated with a food reward (details about task in Supplemental Table 1). Cups containing rewards were covered with digging media, and animals were trained to dig inside the container according to the odor predicting the food, independent of the digging medium covering the cups. After initial training, intradimensional reversals 1 and 2 were introduced, keeping odor as the relevant dimension. Lastly, the extradimensional set-shift phase was initiated with novel stimuli, when the appropriate discriminant was the medium instead of odor. Criterion was set at 6 consecutive correct choices. **B.** Groups did not differ in the total number of trials that animals required to reach criterion or the number of errors made. Arrow indicates that graphs in panel B describe acquisition phase. Box plots represent median, interquartile range and minimum and maximum data values. Detailed statistical analyses are provided in Table 1.

#### Acquisition of the compound stimulus discrimination

Neither AIE exposure nor sex affected the total number of trials that animals required to reach criterion, or the number of errors made. Moreover, no interaction was observed for any parameter (all P’s>0.05). Thus, the process of learning an initial compound discrimination was unaffected by AIE exposure or sex.

#### Reversal 1

From this point, each phase started with the discriminations learned the previous day (reacquisition) before introducing the new rule (reversal; Fig. 3A). No differences were observed in the trials to reach criterion for reacquisition as a function of exposure or sex, and no interaction between factors was identified (all P’s>0.05; Fig. 3B). However, we observed a significant exposure-by-sex interaction on the number of active errors during reacquisition (P=0.05; Fig. 3B). AIE-exposed females exhibited more active errors during reacquisition than controls; however, pairwise differences did not survive Bonferroni correction in post-hoc comparisons.

**Figure 3.**
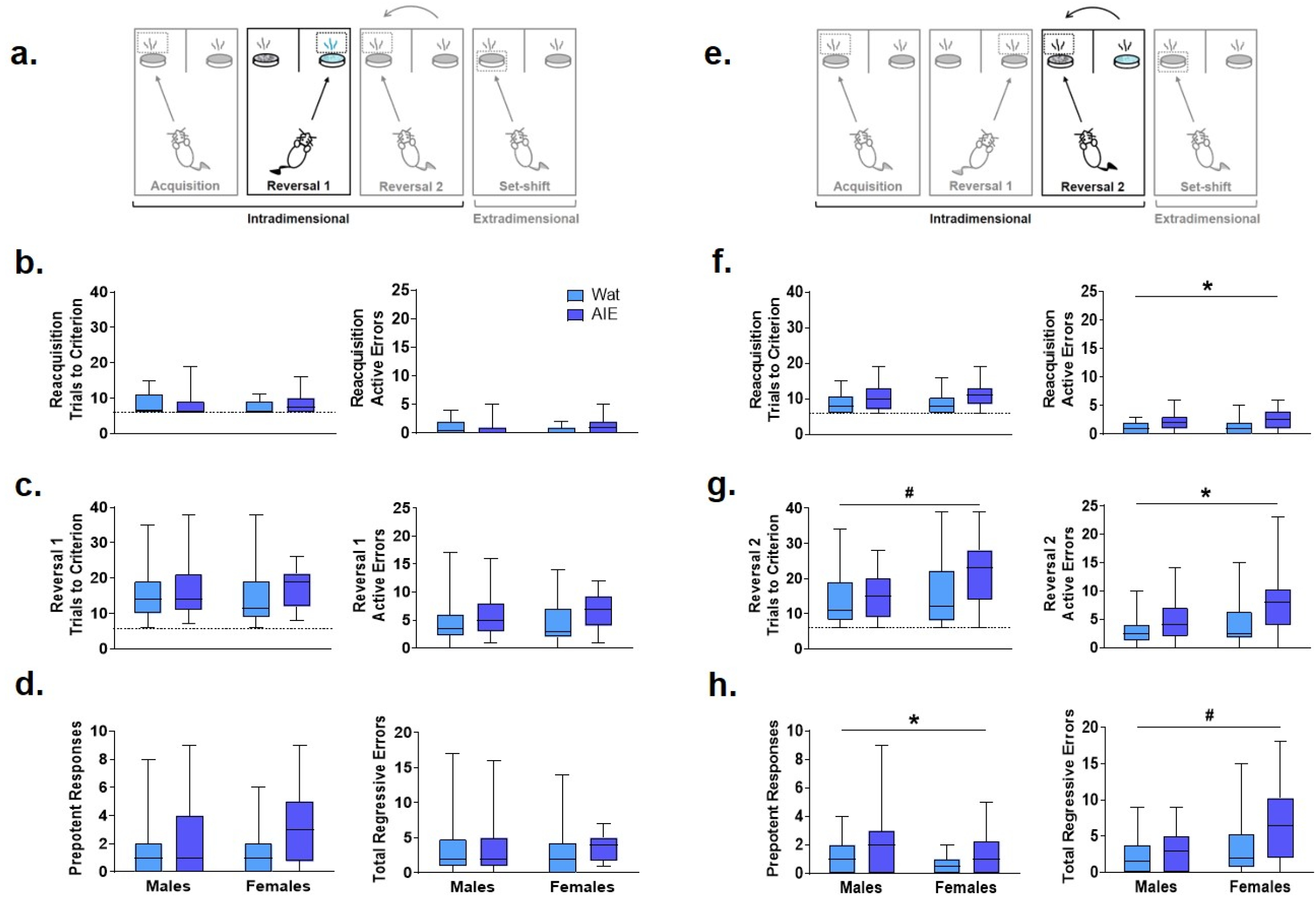
AIE exposure impaired some parameters in reversal learning but not acquisition of rules. **A.** On Reversal 1, the contingency was reversed such that the opposite odor signified the reward location. **B.** Every new training day began with reacquisition of the association from the previous day (reacquisition). Groups did not differ in the number of trials to reach criterion (left). We observed a significant interacting effect of exposure and sex on active errors during reacquisition (right); however, no group differences survived Bonferroni correction. **C.** During Reversal 1, groups did not differ in the total number of trials required to reach criterion (left) or the number of active errors while reaching criterion (right). **D.** As active errors entail different types of errors, we subdivided them into “prepotent” and “regressive,” depending on whether they occurred before or after a correct response, respectively. Groups did not significantly differ in the number of prepotent (left) or regressive (right) errors. **E.** The next day, rats underwent reacquisition followed by Reversal 2, wherein the association was switched back to the original contingency used on Acquisition. **F.** In reacquisition of the previous day’s rule, AIE-exposed animals committed more errors than control rats (right), suggesting that AIE exposure impaired consolidation of what animals learned the day before. **G.** When Reversal 2 was introduced, female rats required more trials than male rats to reach criterion (left). However, AIE-exposed animals committed more total errors than water-exposed rats (right). **H.** Regarding error type, AIE exposure was associated with more prepotent errors (left), while female sex was associated with more regressive errors (right). Statistics and figures for initial regressive and subsequent errors are showed in Supplemental Fig. 2. AIE-exposed animals had more difficulty inhibiting learned rules, updating learned information, and guiding behavioral choices based on feedback than water-exposed animals, effects that were generally larger in females. Box plots show median, interquartile range and minimum and maximum data values. # main effect of sex (P<0.05). * main effect of exposure (P<0.05). Detailed statistical analyses are provided in Table 1.

After rats obtained reacquisition criterion, the contingency of the olfactory cues was switched (Reversal 1). Neither the total number of trials required to reach criterion nor the number of errors committed differed across groups; no main effects of exposure or sex and no exposure-by-sex interaction reached significance (all P’s>0.05; Fig. 3C, Table 1). To better understand the type of errors animals made during their performance, we subdivided errors into “prepotent” and “regressive,” depending on whether they occurred before or after a correct response, respectively. Additionally, regressive errors were split into initial and subsequent regressive errors (see Methods). We observed a trending increase (~50%) in prepotent responses in AIE-exposed animals compared to controls (P=0.08; Fig. 3D), with no main effect of sex or exposure-by-sex interaction (all P’s>0.05, Table 1). As prepotent responses occurred until the first correct response in the new rule, one might expect that positive feedback from reinforcement under the new rule would improve animals’ performance. In fact, that was what we observed during Reversal 1, as regressive errors were similar among groups (Fig. 3D), as were subtypes of regressive errors (Table 1). These results suggest that AIE-exposed animals had some difficulty switching from the previously learned rule as compared to water-exposed siblings. However, after positive feedback, learning of the new rule was similar among groups.

#### Reversal 2

At this stage, both odors had been associated with reward; thus, performance here does not imply new learning. Instead, animals must update their attentional set to a previously learned rule (Fig. 3E). As before, the Reversal 2 phase started with reacquisition of the rule learned the previous day, and AIE-exposed animals needed marginally more trials to reach criterion (P=0.08) and committed more errors (P=0.03; Fig. 3F) than controls. There were no sex differences or exposure-by-sex interactions (all P’s>0.05; Table1), suggesting that AIE exposure impaired consolidation of the contingency the animals learned the day before.

After reacquisition, the original discrimination rule from Day 6 was re-introduced (Reversal 2). AIE- and water-exposed rats did not differ in the number of trials to reach criterion (Table 1); however, female rats required more trials than did males (P=0.04; Fig. 3G). No exposure-by-sex interaction was observed (P>0.05). In analyzing the errors committed during Reversal 2, we found that AIE-exposed animals made more total errors than water-exposed rats (P=0.01; Fig. 3G, Table 1), and female rats showed a marginal increase in total errors compared to males (P=0.10), but no exposure-by-sex interaction was observed (P>0.05). Regarding error type, both AIE exposure (P=0.02) and, marginally, female sex (P=0.06; Fig. 3H, Table 1) were associated with more prepotent errors, but no exposure-by-sex interaction was observed (Table 1). Similarly, total regressive errors were marginally higher in AIE-exposed animals (P=0.10), and significantly higher in female rats (P<0.01; Fig. 3H), with no exposure-by-sex interaction (Table 1). Considering that Reversal 2 involves an update of previous learning, these results indicate that AIE-exposed animals had difficulty either recovering the original rule or keeping track of the current rule, and the positive feedback after a correct response did not maintain performance as it did in Reversal 1.

We further categorized regressive errors as initial and subsequent regressive errors, which reflect distinct deficits: initial regressive errors indicate difficulty in performing a behavior based on the positive feedback after adequate discrimination, while subsequent regressive errors suggest broader deficits in behavior modification based on feedback, whether positive or negative. Initial regressive errors were not different between exposure or sex (all P’s>0.05, Supplemental Fig. 2A, Table 1). However, AIE-exposed animals made more subsequent regressive errors than water-exposed animals (P= 0.05) and females made more subsequent regressive errors than males (P= 0.01, Supplemental Fig. 2B, Table 1). No exposure-by-sex interaction was identified for either initial or subsequent regressive errors (all P’s>0.05). Together, these results suggest that ethanol exposure during adolescence impairs the ability to update information about learned strategies and the capacity to use feedback to guide behavioral choice.

#### Extradimensional Set-Shift

The final phase implemented a new set of digging media and odors along with a change in the relevant sensory dimension, such that the discriminant was now the texture of the medium covering the reward instead of odor (Fig. 4A). Similar to prior testing days, this phase started with reacquisition of the previous day’s rule, where no differences or interactions were observed in the total trials to reach criterion or active errors (all P’s>0.05, Table 1). Next, the extradimensional set-shift began with the new set of stimuli. As in initial acquisition (Day 6), total trials required to reach criterion and active errors were not affected (all P’s>0.05; Fig 4C, Table 1). These results indicate that, despite a change in sensory modality of the relevant stimulus, discrimination learning to novel stimuli was unaltered after AIE exposure, consistent with results observed in the earliest phases of our experiment (Day 6 initial acquisition) and in other studies^9,22^.

**Figure 4.**
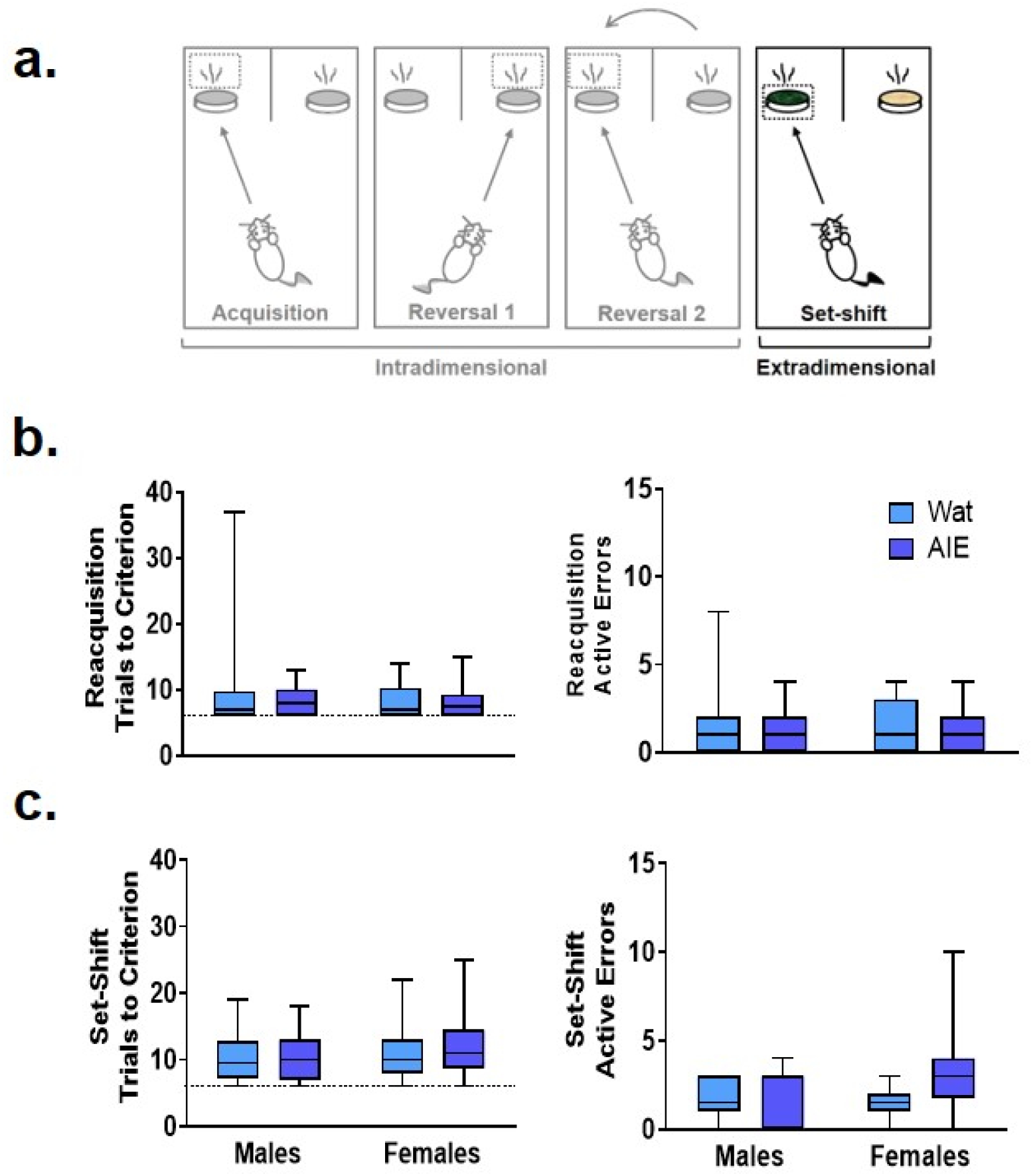
AIE exposure did not impair extradimensional set-shifting performance. **A.** After reversals, an extradimensional set shift to completely new stimuli and sensory modality of the discriminant was introduced. In this phase, the discriminant associated with the reward was now the digging medium and odor was irrelevant. **B.** In the reacquisition phase, groups did not differ in the total trials to reach criterion (left) or active errors (right). **C.** In the extradimensional set shift, groups did not differ in the total trials required to reach criterion (left) or the number of active errors made (right), suggesting that AIE exposure does alter the ability to shift sensory modality of the discriminant in the context of new stimuli. Box plots show median, interquartile range and minimum and maximum data values. Detailed statistical analyses are provided in Table 1.

### Resting state fMRI

We hypothesized that AIE exposure would decrease functional connectivity among regions of interest (ROIs; Fig. 5A) that are known to be involved in action selection and learning: prelimbic cortex (PrL), infralimbic cortex (IL), orbitofrontal cortex (OFC), primary sensory cortex (S1), nucleus accumbens (NAc), caudate-putamen (CPu), dorsal hippocampus (HippD), and thalamus (Thal). To test this, resting-state fMRI was conducted in approximately half of the rats (n=41); behavioral data on this subset of rats is provided in Supplemental Fig. 3 – Fig. 7. We performed a two-way ANOVA followed by post hoc t-tests to determine the effects of sex and AIE exposure on functional connectivity between the pairs of ROI^23^ and used network-based statistics (NBS)^26^ to control the family-wise error rate. AIE exposure significantly reduced functional connectivity (P=0.02), but no main effect of sex or exposure-by-sex interaction was observed (P>0.05). Specifically, AIE exposure reduced functional connectivity between NAc and CPu (T=2.53), NAc and PrL (T=2.54), NAc and S1 (T=2.63), NAc and HippD (T=2.96), and NAc and Thal (T=2.42), as well as IL with HippD (T=2.49), and PrL with Thal (T=2.28) (Fig. 5B). Note that each T-value above indicates that the AIE exposure effect on the connection between two nodes survived the suprathreshold value that set at P < 0.05 for a two-tailed t-test using the NBS approach.

**Figure 5.**
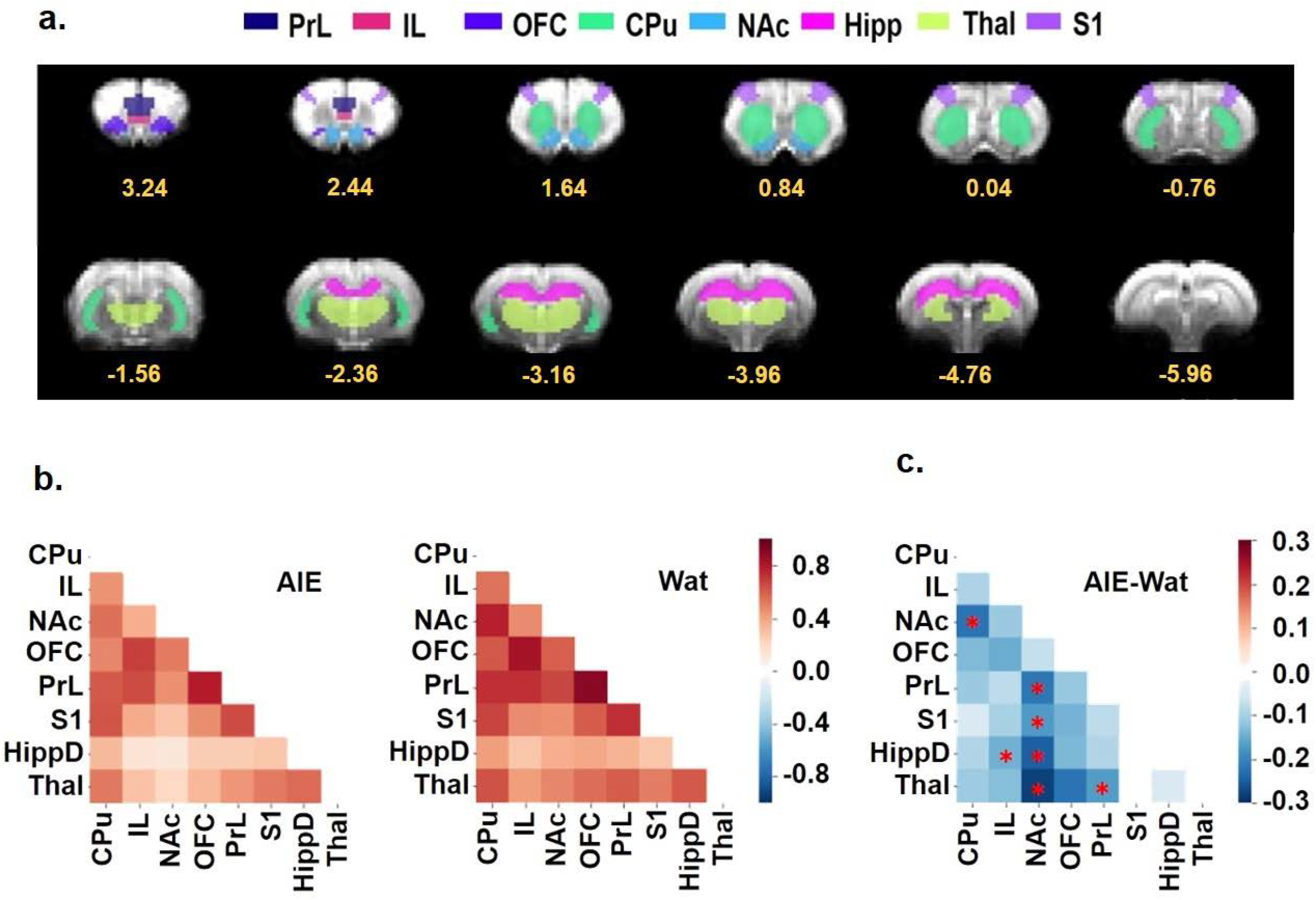
AIE exposure decreased resting-state functional connectivity among regions of interest (ROIs) involved in behavioral flexibility. **A.** ROIs used for analyses shown on the EPI template were manually drawn by co-registering the T2 template to the Paxinos & Watson rat brain atlas^69^, then aligned to the matching EPI templates to ensure that all ROIs were aligned in the correct position within the brain in the fMRI data. Numbers below slices indicate location in mm from bregma. **B.** The Z-score correlation matrixes for the AIE (left) and Wat control (right) groups describe the functional connectivity among ROI pairs, with warm colors (positive Z scores) indicating increased connectivity and cool colors (negative Z scores) indicating decreased connectivity. **C.** AIE-exposed rats exhibited less connectivity among ROI pairs than Wat controls (AIE - Wat, Fisher’s Z transformed, * p<0.05, corrected for family-wise error rate). CPu, caudate-putamen; HippD, hippocampus; IL, infralimbic cortex; NAc, nucleus accumbens, OFC, orbitofrontal cortex; PrL, prelimbic cortex; S1, primary somatosensory cortex; Thal, thalamus; AIE, adolescent intermittent ethanol; Wat: water.

### Correlation between behavior and fMRI

After identifying exposure effects on functional connectivity, we assessed the relationship between functional connectivity among pairs of ROIs and behavior scores, specifically active errors during acquisition of Reversal 2, a behavioral metric that significantly differed by exposure group. To that end, we performed correlation analysis between behavioral data and connectivity values of each ROI pair (Fig. 6A and supplementary Table 2). The results showed that the functional connectivity of CPu-NAc, CPu-OFC, NAc-Thal, and OFC-Thal pairs showed significant negative correlations with the number of active errors during Reversal 2 acquisition (Fig. 6B-6E; all Pearson’s R < −0.35, all P < 0.05). Scatter plots were used to confirm whether this correlation depended on the experimental factors. As observed in Fig. 6B-6E, AIE exposure altered the negative correlation identified in water-exposed rats (right columns) with no effect of sex (Fig. 6, left columns). Together, our results suggest that AIE exposure disrupted the relationship between ROI connectivity and active errors relative to control animals.

**Figure 6.**
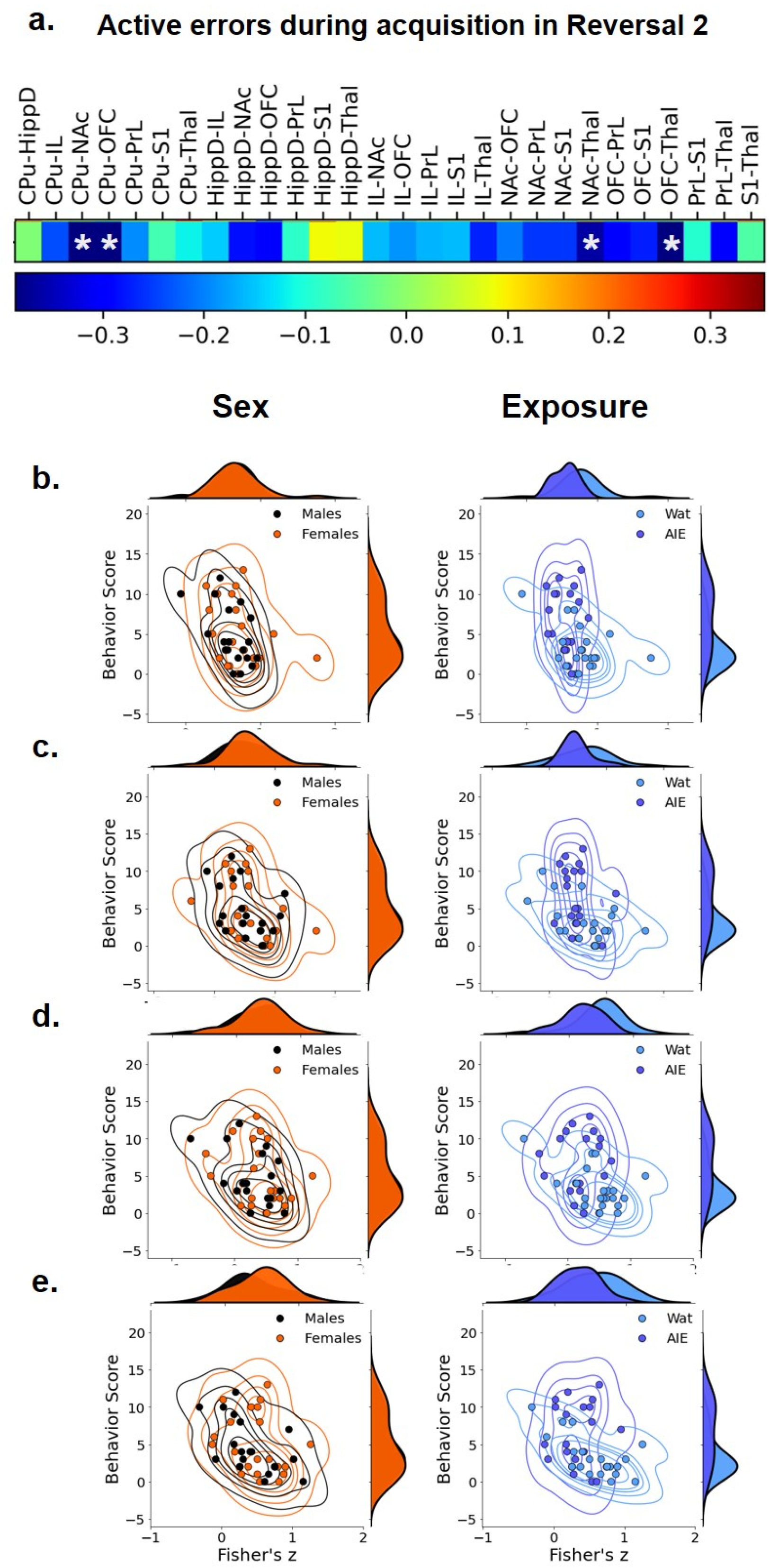
Correlational analysis between active errors during Reversal 2 and functional connectivity among regions of interest (ROIs). We compared the pattern of the values across the subjects between the number of errors in Reversal 2 and pair-wise ROI connectivity to identify the ROI pairs that significantly correlate with behavior at p < 0.05. **A.** In the sequence, we visualized the joint plot for each factor (Sex on left, Exposure on right) to visualize their correlation to the ROI pairs from Figure 5. Most R01 pairs were negatively correlated to active errors in Reversal 2, and four correlations reached statistical significance: **B.** CPu-NAc; **C.** CPu-OFC; **D.** NAc-Thal, and **E.** OFC-Thal. The data suggest that AIE exposure (right correlation plots) affect the behavior/functional connectivity correlation more than sex (left correlation plots). CPu, caudate-putamen; HippD, hippocampus; IL, infralimbic cortex; NAc, nucleus accumbens, OFC, orbitofrontal cortex; PrL, prelimbic cortex; S1, primary somatosensory cortex; Thal, thalamus; AIE, adolescent intermittent ethanol; Wat: water. Detailed correlational analyses are in Supplemental Table 2.

### Mediation analyses

While neuroimaging studies in animal models often have the general goal of generating testable hypotheses for direct interventions (e.g., lesion, chemogenetic, or optogenetic studies), a second important use of such studies is for translational comparisons, especially to data from human subjects. Primarily in support of the latter goal, we next tested whether AIE exposure effects on brain connectivity mediated AIE exposure effects on performance in the attentional set-shift task. To do so, we conducted mediation analyses (Fig. 7), a statistical method to assess whether inclusion of a mediating variable *(M)* significantly alters the slope of the relationship between two other variables *(X* and *Y)*^27^. In this case, *X* represents AIE exposure, *Y* represents set-shift task performance, and *M* variables were measures of pairwise connectivity between ROI. Due to the large number of pairwise brain connections that could serve as potential mediators, the high correlation among ROIs, and the consistent negative direction of AIE exposure effects on brain connectivity, we used a dimensionality reduction approach to calculate a single measure of brain functional connectivity. First, we selected an informative set of pairwise connections which varied with AIE exposure at a lenient statistical threshold of *p*<0.1, uncorrected (*n*=11; Fig. 7A). This somewhat lenient threshold was chosen since connections with weaker associations in univariate analyses can still contribute meaningful variance when combined with other variables^28^. Next, we conducted a principal component analysis of this reduced functional connectivity matrix, which yielded a first principal component accounting for 66% of the variance in functional connectivity among the included connections. This principal component reflects a “subnetwork” of functional connections that were most affected by AIE exposure (Fig. 7A), and each connection contributed a different weight to the principal component-derived subnetwork (Fig. 7B). Of note, this subnetwork included three of the four connections that were independently associated with errors during Reversal 2 based on univariate correlations (i.e., CPu-NAc, NAc-Thal, and OFC-Thal; Figure 6). Next, we assessed whether AIE exposure effects on this functional subnetwork mediated the effects of AIE exposure on the number of errors committed during Reversal 2. The analysis indicated that AIE exposure effects on active errors during Reversal 2 were significantly mediated by brain functional connectivity (P<0.001; Fig. 7C). Moreover, functional connectivity in the principal component subnetwork accounted for 32% of the total effect of AIE exposure on active errors during Reversal 2 (P=0.03). Thus, AIE-induced reductions in functional connectivity across brain regions involved in action selection and learning meaningfully contributed to the observed behavioral flexibility deficits.

**Figure 7.**
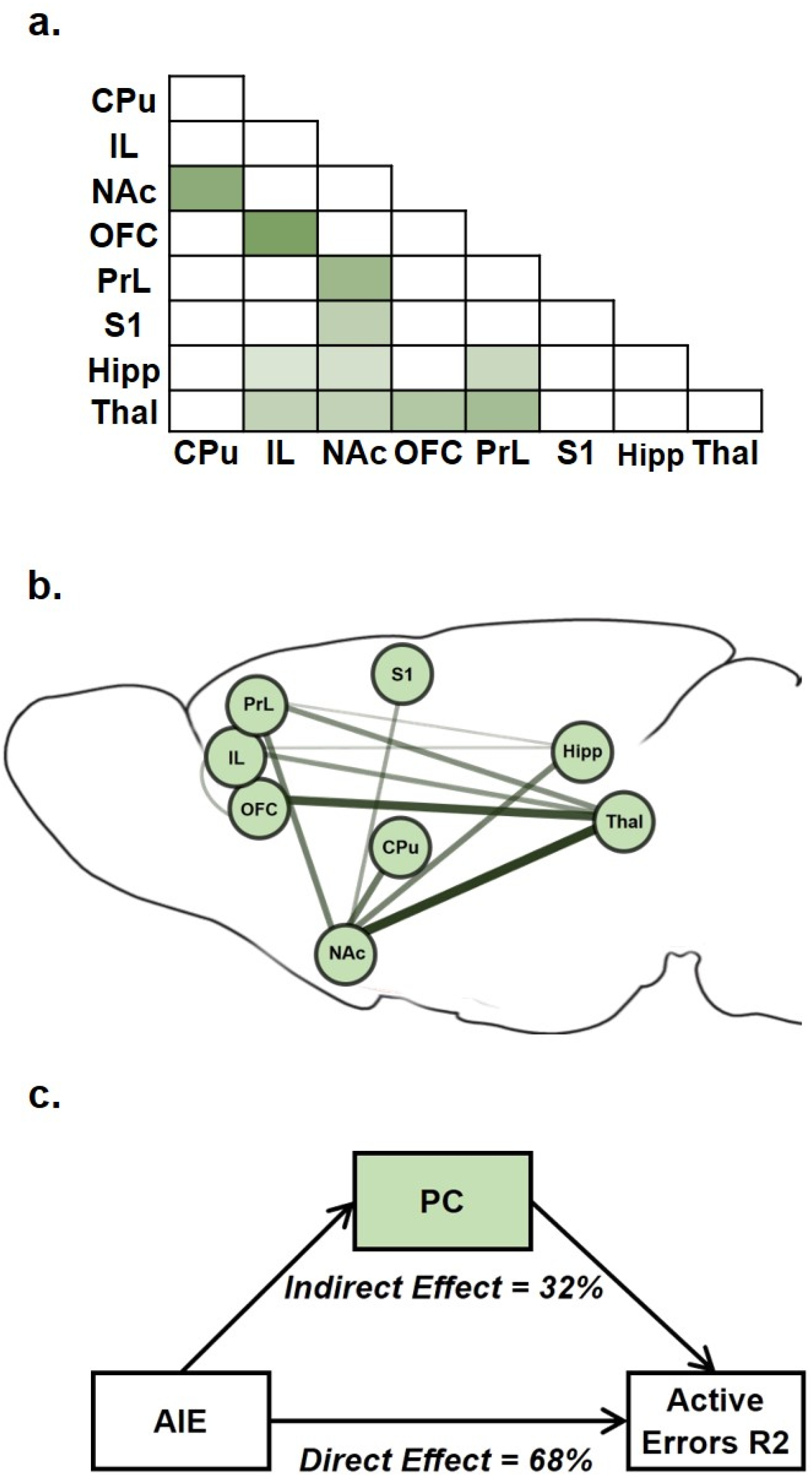
AIE-related changes in brain functional connectivity mediated AIE exposure effects on active errors during Reversal 2. **A.** Average correlation matrix representing functional connectivity between regions-of-interest (ROIs) for the 11 connections which demonstrated an effect of exposure (p<0.1) and were included in the subsequent subnetwork analysis. Darker color corresponds to stronger functional connectivity. **B.** A singular value decomposition produced a principal component that explained 66% of shared variance among the 11 connections. The subnetwork identified by the principal component is depicted, with the size and darkness of lines connecting ROIs pairs indicating component weights. **C.** In the mediation analysis, the principal component subnetwork of functional connections accounted for 32% of the effect of exposure on behavior that was explained by the indirect effect through network functional connectivity is indicated. CPu, caudate-putamen; HippD, hippocampus; IL, infralimbic cortex; NAc, nucleus accumbens, OFC, orbitofrontal cortex; PrL, prelimbic cortex; S1, primary somatosensory cortex; Thal, thalamus; AIE, adolescent intermittent ethanol.

## Discussion

Behavioral flexibility is a complex construct requiring several constituent functions that recruit multiple brain circuits. While prior studies reported that AIE exposure induces deficits in behavioral flexibility in adult rodents^9,18,19,22^, the behavioral deficits could not be linked to changes in brain circuit function. In this study, we investigated AIE exposure effects on conditioned responding to reward-associated cues (Pavlovian conditioned approach), flexible decision-making (attentional set-shifting task), and resting-state functional connectivity MRI in the same animals to allow direct linkage of functional connectivity among brain regions involved in action selection to behavioral deficits. While we did not observe AIE exposure effects on conditioned approach in this study, we did observe clear deficits in reversal learning and a marked reduction in functional connectivity among ROI. We also found a clear association between functional connectivity measures and behavioral performance. Moreover, we identified a functional connectivity subnetwork that mediated reversal learning deficits, demonstrating statistical causation between the degree of functional connectivity and behavioral flexibility. These complimentary analyses identified especially strong contributions of the NAc, thalamus, and OFC in the AIE-induced deficits in behavioral flexibility, a circuit associated with compulsive and inflexible behavior in substance use disorders^29^.

Behaviorally, we observed robust effects of AIE exposure on attentional set-shifting performance. Phases implying new learning (e.g., initial acquisition) were not affected by AIE exposure, consistent with previous findings^9,30^, showing that skills required to associate and discriminate stimuli are spared after AIE exposure. However, reacquisition was affected by AIE exposure, even though rats achieved the performance criterion the day before, suggesting deficits in consolidation. Before Reversal 1, AIE exposure impacted reacquisition differently by sex, with AIE-exposed females exhibiting more errors than control females, although the post-hoc comparisons did not reach significance. We also observed AIE-induced deficits in both males and females during reacquisition before Reversal 2, indicating that although the new rule of Reversal 1 was learned, it was not as effectively consolidated among AIE-exposed animals compared to controls. As AIE-induced impairments in reacquisition performance were more robust before Reversal 2 (that assessed the reversed contingency) than before Reversal 1 (that assessed the original contingency), it is possible that learning of the original contingency was better consolidated in AIE-exposed rats than the reversed contingency. This interpretation is supported by the finding that AIE exposure did not alter reacquisition before the Extradimensional Set-Shift, which also assessed the original contingency. These results are only in partial agreement with our prior study^22^, in which AIE exposure did not significantly impact any reacquisition phase. Reacquisition is not typically reported^9,16,17^, and we are not aware of studies evaluating the effect of AIE exposure on memory consolidation. However, AIE exposure alters adolescent hippocampal development^31,32^, a critical region involved in memory consolidation through “hippocampal–neocortical dialogue” (see^33^ for review).

The introduction of reversal phases revealed interesting differences among AIE-exposed and control groups. In Reversal 1, we observed no differences regarding total trials to criterion or errors, although prepotent responses were marginally different in that AIE-exposed subjects required more trials to adapt to the new rule. In Reversal 2, AIE-exposed rats made more active errors, and specifically more prepotent responses and subsequent regressive errors, while learning the new rule. Thus, AIE-exposed rats were slower to incorporate negative feedback and initiate a new strategy^34^ in both reversals, but this effect was stronger in Reversal 2. In contrast, in our previous study^22^, AIE-induced deficits were observed in Reversal 1, while AIE-exposed rats performed better in Reversal 2. This pattern of behavior indicated that the original attentional set dominated behavioral choice, slowing the ability to learn an initial reversal but facilitating return to that contingency in Reversal 2. A major difference is that in the present study, rats had Pavlovian conditioning training prior to assessing attentional set learning, and previous behavioral training can impact subsequent task performance^16,35^. However, both studies identified increased prepotent responses in the AIE-exposed groups, consistent with perseveration on the previous response despite negative feedback. This particular deficit is suggestive of impaired inhibitory control^1,2^ and is consistent with other studies using AIE^16–18^ and chronic intermittent ethanol (CIE) exposure in adult rodents^36^, and even in humans (for a recent review see^37^). Moreover, the present study found that subsequent regressive errors were higher after AIE exposure. In general, regressive errors suggest a lack of ability to maintain a new association or keep the current rule “in mind” despite feedback. Thus, our results suggest an AIE-induced deficit in working memory, a conclusion supported by other studies showing adolescent alcohol effects on that function in animals^12,13^ and, less consistently, in humans^38,39^. Overall, these reversal data suggest that AIE exposure induces deficits in the fundamental processes of inhibitory control and working memory, resulting in deficits in flexible decision-making. Future studies can directly assess inhibitory control and working memory in animals with and without AIE exposure to confirm this interpretation.

Perseveration of an acquired attentional set illustrates how well-learned cues can retain control over action selection after AIE exposure. However, perseveration did not extend to the sensory domain of the relevant cue: when novel odor and tactile stimuli were used in the extradimensional set shift, all rats readily learned to pay attention to the tactile rather than the odor cue, consistent with previous studies^22,36^ using the digging task to assess alcohol effects on attentional sets. In contrast, studies using an operant-chamber version of a set-shift from a visual rule to a location rule reported AIE-induced deficits after chronic alcohol exposure^40^ and AIE^9,16^ (with one study only observing an AIE effect in males^41^). However, in this latter task, the formerly relevant cues remain in the chamber, and the subject must inhibit responses to the visual cues and respond instead by location. Thus, those data can also be interpreted as an inability to inhibit a previously learned attentional set in AIE-exposed animals, as opposed to inability to shift the relevant stimulus from one domain to another.

While we replicated a sex difference in sign-tracking behavior in Pavlovian conditioned approach^19^, we did not observe an effect of AIE exposure on conditioned approach in the present study, in contrast to other reports in the literature^14,15,19^. Large individual differences are often observed in conditioned approach due to considerable variability among animals in the propensity to attribute incentive salience to cues^42,43^. In addition, there is large variability in the field around Pavlovian conditioning methodology, such as the timing of the cue and the length of training, and these factors may have contributed to our outcome. While we did not replicate the previous findings, indicating that the effect of AIE exposure on conditioned responses may be less robust than previously observed, it remains that the majority of studies^14,15,19^ to date have reported shifts toward sign-tracking after AIE exposure, suggesting enhanced sensitivity to reward-associated cues or incentive salience attribution. It’s worth noting that we did not test the persistence of conditioned approach behavior (e.g., via reward omission) or the development of incentive salience of the conditioned cue (e.g., via Pavlovian instrumental transfer). Thus, an empirical question is whether flexibility in these aspects of reward conditioning is altered by AIE exposure.

Action selection and behavioral flexibility in response to changing circumstances are largely mediated by cortico-striatal-thalamic circuits^44^. Moreover, gaps in memory encoding or consolidation might be associated with reduced function of prefrontal and hippocampal areas^45,46^, deficits in inhibitory control with fronto-striatal regions^1,37^, and impairments in working memory and behavioral flexibility mainly with prefrontal regions^1,6^ – all brain regions that have been shown to be affected by AIE exposure^6^. Thus, we hypothesized that behavioral deficits would be associated with altered functional connectivity among fronto-limbic regions in AIE-exposed animals. We found that AIE exposure reduced functional connectivity across multiple ROI pairs, including frontal, hippocampal and thalamic connections with the NAc. We previously observed similar AIE-induced reductions in functional connectivity across fronto-striatal regions of male rats^23^, and the present study extended those findings to females. Linking the MRI data to behavioral assessments, the correlational analysis between pairwise connectivity strength and behavioral performance in Reversal 2 showed a negative association between functional connectivity in specific ROI pairs (including OFC, NAc, CPu, and Thal) and observed behavioral deficits. Interestingly, the correlation was stronger in water-exposed rats and reduced in AIE-exposed rats, suggesting that ethanol disrupted the association between functional connectivity and the behavior. Since we were underpowered to statistically validate this statement, additional studies including more animals can clarify this effect. An additional aspect derived from this analysis was the alleged key role that connections between OFC, NAc, and Thal play in the effects of AIE exposure on behavioral deficits, something also observed in the mediation analysis.

To address the interconnected nature of the ROIs, we used principal component analysis of functional connections to identify a principal component or “subnetwork” that might mediate observed behavioral deficits. The functional connections of the identified subnetwork (accounting for 66% of variance) include brain regions and projections observed in several studies to be involved in the cognitive processes of flexible decision-making^2,47–54^. Interestingly, two connections that contributed most strongly to this subnetwork – NAc-Thal and Thal-OFC – form the striato-thalamo-orbitofrontal circuit that is hypothesized to be persistently impaired after repeated exposure to drugs of abuse and to contribute to the compulsive and inflexible behaviors associated with addiction^29^. The finding that adolescent alcohol exposure, in the absence of “addiction,” can affect similar circuits as those affected by chronic substance use supports an apparent vulnerability of prefrontal structures implicated in “top-down” subcortical regulation to drug exposure^6,7,29^. Using this subnetwork as a mediator, we determined that AIE-associated decreases in functional connectivity mediated the increases in active errors during Reversal 2, when both contingencies had been experienced. While at least one prior study has correlated specific MRI measures to animal behavior^55^, to our knowledge, this is the first study to demonstrate that changes in functional connectivity statistically mediated behavioral performance in the same set of animals. Thus, it is validating that the different approaches to assess the impact of AIE exposure on brain functional connectivity yielded converging results highlighting connections between prefrontal, accumbal and thalamic regions.

There are many possible mechanisms that may underlie this altered functional connectivity following adolescent alcohol exposure, one of which is AIE-induced changes in cholinergic function. There is compelling evidence showing decreases in forebrain cholinergic phenotypic markers^16–18,56^ and receptors^18^ after AIE exposure in rodents, supporting AIE-induced reductions in cholinergic neurotransmission that persist into adulthood and likely affect behavior (see^6^ for a review). Basal forebrain cholinergic neurons project extensively throughout the brain, including several regions in the functional connectivity subnetwork we identified: prefrontal cortex, hippocampus, and sensory cortex receive direct, modulatory cholinergic projections from the basal forebrain^57,58^. Documented cholinergic involvement in cognitive processes like learning, attention and memory (including working memory)^59,60^ implicates acetylcholine as a crucial player in cognitive performance. Thus, changes in cholinergic dynamics might explain behavioral and connectivity impairments in AIE-exposed animals; however, this hypothesis remains to be directly tested.

A strength of this study is large number of animals included, particularly for the behavioral measurements, which allowed for assessment of sex as a biological variable. Although no statistical differences were observed in functional connectivity among males and females, we found that females displayed a stronger deficit in reversal learning after AIE exposure and more sign-tracking behavior relative to males. While a sex difference in sign-tracking behavior is consistent with the literature^22,61,62^, many studies assessing reversal learning after AIE exposure have only investigated cognitive deficits in males^9^, were not sufficiently powered to address sex differences^22^, or presented similar AIE effects between males and females^17^. However, understanding the underlying mechanisms that are sex-specific, such as immune reactivity and maturation of neural networks^63^, may explain the domain, severity and persistence of cognitive deficits in females versus males. A limitation of the present study is that animals were sedated during MRI scans rather than awake. While awake rat fMRI is possible, we used the most widely used light anesthesia protocol^64^ to avoid the potential confounds of habituation and restraint after our behavioral approaches. Importantly, studies have shown that major brain connectivity changes are preserved across anesthesia states^65^.

In summary, we identified a functional connectivity subnetwork based on selected ROIs that was highly predictive of behavioral performance and mediated the effect of AIE exposure on reversal learning. This also suggests an effect of adolescent alcohol exposure on the striato-thalamo-orbitofrontal circuit, which has been related to compulsive behavior and risk for developing substance use disorders^29^. Here, we provided support for the hypothesis that decreased functional connectivity after AIE exposure mediates deficits in cognitive processing that impairs flexible decision-making.

## Methods

### Subjects

In this study, we included seventy-nine Sprague-Dawley rats (male n=39, and female n=40), bred and raised in-house. Vivarium temperature and humidity were kept constant with a 12:12 hour light cycle applied (lights on at 0700). After birth (postnatal day [P] 1), litters were culled to 8-10 pups with a similar proportion of males and females. As in previous studies, animals were weaned on P21 and pair-housed with a same-sex littermate throughout all testing. Rats received *ad libitum* food and water access with the exception of food-restriction (85% of their free-feeding body weight) beginning 14 days prior to the attentional set-shift task until the fMRI scan was complete. All experiments were performed following the NIH Guide for the Care and Use of Laboratory Animals with procedures approved by the Institutional Animal Care and Use Committee of the University of North Carolina at Chapel Hill.

### Adolescent Intermittent Ethanol Exposure

Starting at P25, animals were weighed and given 5 g/kg of ethanol (25% v/v in water) or an equivalent volume of water (controls) intragastrically (i.g.), in an intermittent pattern (once/day on a 2 days on, 2 days off schedule) until P54 for a total of 16 exposures (Fig 1A). This dose produces blood alcohol levels of approximately 230 mg/dl in males and females 60 min after administration^19^. Animals housed together were given the same exposure (either alcohol or water) throughout adolescence.

### Pavlovian Conditioned Approach

Operant chambers (Med Associates, St. Albans, VT) were used to train animals in Pavlovian conditioned approach^19^. At ~P70, rats were given home-cage access to 20% sucrose in water for 1 hour prior to magazine training to become familiarized with the reward. After that, animals were placed in behavioral chambers (12” length × 12.5” width × 11.5” height) for receptacle training to familiarize them with sucrose delivery every 120 to 230 seconds into the receptacle. After the session, the receptacle was checked to verify that the animal consumed the reward. Next, Pavlovian conditioning occurred daily for 20 days across 4 weeks (Fig. 1A). Rats were presented with a conditioned stimulus (CS; illuminated cue light plus lever extension) for 30 seconds. At the end of the 30-second period, the lever retracted, the light turned off, and a reward (100 μl 20% sucrose in water) was delivered into the receptacle. Fifteen trials occurred throughout the session with a variable inter-trial interval (90 to 210 seconds). Behavioral measures (lever presses, receptacle entries) and conditioning events (CS+ onset and offset, sucrose delivery) were recorded and timestamped by Med Associates for later analysis.

We monitored both lever presses and receptacle entries, and calculated the latency to press the lever or enter the receptacle after the onset of the CS. The probability to press the lever or enter the receptacle during the CS was measured as the number of trials that each behavior occurred divided by 15 (the total number of trials). For receptacle entries, we calculated an elevation score, measured as the number of receptacle entries during a CS presentation less the number of receptacle entries 30 seconds prior; this described those entries that were conditioned responses to the CS^24^.

### Attentional set-shift task

Discriminative association was measured using a digging task^22,25^, wherein a reward (1/4 of a fruit loop cereal) was buried in digging media and associated with a specific odor (e.g., vanilla) placed in one of two dishes (see Fig. 2A and Supplemental Table 1 for a detailed description). Animals were required to perform six correct choices in a row to reach criterion and advance to the next step.

On training days 1-3, rats learned to retrieve a reward from one of two dishes at the end of the chamber unencumbered by digging media at first. On days 4-5, dishes contained digging media (white or brown bedding) and the rat learned to dig to find and consume the reward.

Acquisition of the discriminative attentional set began on day 6. Randomizing the digging media and side of the box, the reward was placed in a dish with a corresponding odor (e.g., vanilla) while the other dish (e.g., cinnamon) did not contain a reward. On each trial, the rat chose a dish by digging with the face or paws; at that point, entry to the other dish was blocked. If correct, the rat was given time to consume the reward. If incorrect, the rat was given time to realize no reward was present. If the rat did not make a choice within a 3-minute period, the trial was recorded as an omission and counted against their criterion. When the rat chose the correct dish six consecutive times, the rat reached criterion and finished testing for that day.

On the first reversal day (Reversal 1, day 7; Fig. 3A), the discrimination task began with reacquisition of the association from the previous day. Once the rat reached criterion for reacquisition (6 consecutive correct choices), the contingency was reversed such that the opposite odor (e.g., cinnamon) signified the reward location. Rats continued testing until criterion was met.

A second reversal was conducted the next day (Reversal 2, day 8; Fig. 3E) wherein the association was switched back to the original contingency used on day 6 after reacquisition was completed. Rats once again continued testing until criterion was met and data for all trials were recorded.

The following day (Extradimensional Set-Shift, day 9, Fig. 4A), after the rat completed reacquisition, there was an extradimensional set-shift of sensory modality of the discriminant using novel stimuli. If vanilla and cinnamon were the odors previously used, the set-shift used coconut and paprika. Likewise, if white and brown crinkle paper were previously used, the set-shift used gravel and sand. Importantly, the discriminant associated with the reward was now the digging medium and odor was irrelevant.

Errors after a contingency reversal were categorized as prepotent or regressive. Upon reversal, errors made prior to a correct choice within the new contingency were recorded as prepotent responses. Once a correct choice had been made, and a reward was reached (positive feedback), all future errors in that contingency were recorded as regressive errors. Regressive errors were further divided according to whether it occurred after a correct choice (initial regressive error) or after an error itself (subsequent regressive error). As mentioned in the results, initial and subsequent regressive errors reflect distinct deficits: initial regressive errors indicate difficulties in performing a behavior based on the positive feedback after adequate discrimination, while subsequent regressive errors suggest deficits in behavior modification based on feedback, whether positive or negative. This difference provides information on how much the cues and the presence or absence of the reward control behavior, as well as the cognitive processes required to maintain information about relevant contingencies.

### Resting state fMRI

#### Animal Preparation

The MRI study was performed on a subset of 41 rats (male n=19, and female n=22) from the larger sample that underwent the Pavlovian conditioned approach and set-shifting task. The endotracheal tube (Surflash Polyurethane IV Catheter 14G x 2”, TERUMO, Somerset, NJ, USA) was intubated immediately after anesthetic induction with 4% isoflurane to conduct mechanical ventilation at 60 breaths/min with an inspiration time at 40% ratio using MRI-Compatible Ventilator (MRI-1, CWE Inc., Ardmore, PA, USA). The animals were anesthetized under 2% isoflurane (Isoflurane Vaporizer #911103, VetEquip Inc., Livermore, CA, USA) during positioning on a custom-built cradle. MR compatible sensors were installed to monitor core body temperature (OAKTON Temp9500, Cole-Parmer, Vernon Hills, IL, USA), heart rate, peripheral blood oxygen saturation, and end-tidal CO_2_ (SURGIVET^®^ V90041LF, Smith Medical, Dublin, OH, USA). The body temperature was maintained at 37 ± 0.5°C using a circulating water blanket connected to a temperature adjustable water bath (Haake S13, Thermo Fisher Scientific, Waltham, MA, USA). The tidal volume of mechanical ventilation was adjusted to keep the heart rate at 300 ± 50 beats per minute, peripheral blood oxygen saturation above 96%, and end-tidal CO_2_ between 2.8 - 3.2%. To reliably probe functional connectivity, a continuous infusion of dexmedetomidine (0.05 mg/kg/hr) and pancuronium bromide (0.5 mg/kg/hr) was initiated 30 min before the MRI scans and isoflurane was reduced to 0.5% before fMRI experiments^23^.

#### fMRI acquisition

All MR images were conducted on a Bruker BioSpec 9.4-Tesla, 30 cm bore system. A 72 mm volume transmitter coil and a 4-channel receiver array coil were used. Magnetic field homogeneity was optimized first by global shimming, followed by local second-order shims using a MAPSHIM map protocol. We acquired a single 30 min BOLD fMRI scan using a 2D multi-slice, single-shot, gradient-echo, echo-planar imaging (EPI) sequence [scan parameters: TR (repetition time) = 2000 ms, TE (echo time) = 14 ms, bandwidth = 250kHz, flip angle = 70 degrees, voxel size = 0.4 x 0.4 x 0.4 mm^3^, matrix size = 72 x 72 x 32, and FOV (field of view) = 28.8 x 28.8 x 12.8 mm^3^].

#### Data preprocessing

Raw scan data were downloaded from the Bruker 9.4T scanner, converted into NifTI-1 data format, and organized based on the Brain Imaging Data Structure (BIDS) guideline using an in-house built converter. The data preprocessing was initiated with the slice timing correction and rigid body motion correction using the command-line interfaces of Analysis of Functional Neuroimages (AFNI) software^66^. Manual skull stripping was performed on the first frame of the functional connectivity MRI image to precisely employ non-linear spatial normalization using the Symmetric Normalization (SyN) algorithm of the advanced normalization tools (ANTs) package^67^. The EPI-based brain template was used as a reference for spatial normalization. Finally, we performed signal processing on a time-domain signal using SciPy^68^ python library. These signal processes include nuisance signal regression with six rigid motion parameters, bandpass filtering at 0.01 - 0.15 Hz, and spatial smoothing using a Gaussian kernel function with a full width at half maximum (FWHM) expression extent at 0.5 mm. Our automated preprocessing pipeline was built using Python Neuroimaging Pipeline Tool (PyNIPT), an open-source pipeline framework, and available at GitHub repository (https://github.com/CAMRIatUNC).

### Data Analysis

#### Behavior

Statistical analyses of behavioral data were conducted using SPSS for Microsoft (version 26, available from the Virtual Lab of the University of North Carolina at Chapel Hill). Most behavioral data were not normally distributed; thus, the effects of AIE exposure and sex on behavioral metrics from conditioned approach and the attentional set-shifting task were analyzed using a generalized linear model with a Poisson distribution and a log link function. Post-hoc comparisons were Bonferroni-corrected. The elevation score (normally distributed) was analyzed using a two-way ANOVA in GraphPad (version 8, San Diego, CA). Statistical significance was set at p ≤ 0.05 and marginal significance was set at p ≤ 0.10.

#### MRI ROI analysis

We determined eight ROIs (Fig. 5A): the prelimbic prefrontal cortex (PrL), infralimbic cortex (IL), nucleus accumbens (NAc), caudate putamen (CPu), orbitofrontal cortex (OFC), dorsal hippocampus (HippD), thalamus (Thal), and primary somatosensory cortex (S1). All ROIs were manually drawn on a T2-weighted MRI template co-registered to the Paxinos & Watson rat brain atlas^69^ (Fig. 5A), then warped to the matching EPI MRI templates to ensure that all ROIs were aligned in the correct position within the brain in the fMRI data. The resting-state BOLD signal in each ROI was extracted from the preprocessed EPI data that had been aligned to this EPI MRI template and averaged to use for correlation analysis. Pearson’s correlation coefficients were computed in pairs between ROIs to create a connectivity matrix at the individual level. Fisher’s transform was applied to convert the correlation coefficient into approximately normal distribution (Fisher’s z score) for group statistics. Two-way ANOVA was conducted with 2 × 2 between-subject (sex and alcohol exposure) design using statsmodels^70^, a Python module for statistical modeling. To identify significant changes in connectivity, the suprathreshold of p < 0.05 was applied to build sparse matrix for each subject, and we controlled for multiple comparisons by employing link-based family-wise error rate using the network-based statistics^26^ approach. Significant changes in connectivity in pairwise ROIs were determined by thresholding at a corrected p-value < 0.05.

#### Correlational Analyses

To assess the statistical correlation between behavior and fMRI, we calculated the Pearson’s correlation coefficient between behavior scores and the functional connectivity for each pair of regions. The p values were estimated based on the beta distribution on the interval from −1 to 1 with equal shape parameters a = b = n/2 – 1. Subsequently, a false discovery rate correction was applied to control for multiple comparisons. Like the mediational analysis, we focused on active errors during acquisition in Reversal 2 as these showed robust behavioral effects of AIE exposure. Statistical significance was set at a corrected p < 0.05.

#### Mediation Analyses

To test the hypothesis that the effects of AIE exposure on behavior (i.e., AIE → behavior) were mediated by AIE-related changes in brain functional connectivity (i.e., AIE → brain functional connectivity → behavior), we conducted a causal mediation analysis. Causal mediation is a method of assessing the directed or causal relationships among variables^71^. Due to the large number of brain connections that could serve as potential mediators, as well as the numerous positive correlations among the ROIs, we took a dimensionality-reduction approach to examine brain functional connectivity effects at the subnetwork-level, rather than test individual ROI-pairs. We employed a singular value decomposition to obtain the first principal component of the brain functional connectivity matrix, limiting the analysis to those connections (n=11) for which the effect of AIE exposure was p < 0.1, uncorrected (Fig. 7A). The resultant component explained 66% of the variance among the 11 included connections, and the positive weights for each connection were consistent with the interpretation of the set of connections as a cohesive functional subnetwork of the included ROIs (Fig. 7A). The component score for each subject was entered as the brain functional connectivity mediator in the mediation analysis (Fig. 7C).

Mediation analyses were conducted in SAS 9.4 using the CAUSALMED Procedure^71^ using a counterfactual framework^72^, which allows for testing of nonlinear effects such as count data, for active errors during acquisition in Reversal 2 that demonstrated consistent AIE exposure effects in both the MRI subsample and the larger sample (see Table 1). The effect of AIE exposure on brain functional connectivity was modeled with a normal distribution, whereas the effects of AIE exposure and brain functional connectivity on the behavioral variables were modeled with a Poisson distribution using a log link function. Sex was included as a covariate.

## Acknowledgements

The authors thank Dr. Margaret Broadwater of the Biomedical Research Imaging Center in the University of North Carolina at Chapel Hill for her valuable support during the initial phases of this project.

## Author Contributions

DLR and CAB designed the studies; AGA, CAD, AE, SHL and WB collected and analyzed the data; AGA, AE, SHL, YIS, CAB and DLR interpreted the data; AGA, CAD, AE, SHL, CAB and DLR wrote the manuscript; all authors edited and approved the manuscript.

