## Supplemental Data for "Altered cortico-subcortical network after adolescent alcohol exposure mediates behavioral deficits in flexible decision-making"

### Supplemental Material

Alexander Gómez-A[1], Carol A. Dannenhoffer[1], Amanda Elton[2,3], SungHo Lee[3,4], Woomi Ban[3], Yen-Yu Ian Shih[1,3,4,5], Charlotte A. Boettiger[1,2,3,5] & Donita L. Robinson[1,5,6]*

[1] Bowles Center for Alcohol Studies University of North Carolina, Chapel Hill, NC, USA

[2] Department of Psychology and Neuroscience, University of North Carolina, Chapel Hill, NC, USA

[3] Biomedical Research Imaging Center, University of North Carolina, Chapel Hill, NC USA

[4] Department of Neurology, University of North Carolina, Chapel Hill, NC, USA

[5] Neuroscience Curriculum, University of North Carolina, Chapel Hill, NC, USA

[6] Department of Psychiatry, University of North Carolina, Chapel Hill, NC, USA

* Corresponding author

Author information: Dr. Donita L. Robinson, Bowles Center for Alcohol Studies, CB 7178, University of North Carolina, NC 27599-7178, USA, 1+ 919-966-9178

**Supplemental Table 1:** Behavioral training and description of the attentional set-shift task.

| Day | Task | Performance Criterion | Dimensions | | Example combinations | |
|---|---|---|---|---|---|---|
| | | | *Relevant* | *Irrelevant* | *Correct Response* | *Error* |
| 1-3 | Habituation without media | 15-min session per day; 3 min per trial | | | | |
| 4-5 | Habituation with media | Retrieve 20 rewards | | | | |
| 6 | Compound discrimination | 6 consecutive correct trials | Odor | Medium | Vanilla/white paper squares Vanilla/brown paper bedding | Cinnamon/white paper squares Cinnamon/brown paper bedding |
| 7 | Reacquisition | 6 consecutive correct trials | Odor | Medium | Vanilla/white paper squares Vanilla/brown paper bedding | Cinnamon/white paper squares Cinnamon/brown paper bedding |
| | Reversal 1 | 6 consecutive correct trials | Odor | Medium | Cinnamon/white paper squares Cinnamon/brown paper bedding | Vanilla/white paper squares Vanilla/brown paper bedding |
| 8 | Reacquisition | 6 consecutive correct trials | Odor | Medium | Cinnamon/white paper squares Cinnamon/brown paper bedding | Vanilla/white paper squares Vanilla/brown paper bedding |
| | Reversal 2 | 6 consecutive correct trials | Odor | Medium | Vanilla/white paper squares Vanilla/brown paper bedding | Cinnamon/white paper squares Cinnamon/brown paper bedding |
| 9 | Reacquisition | 6 consecutive correct trials | Odor | Medium | Vanilla/white paper squares Vanilla/brown paper bedding | Cinnamon/white paper squares Cinnamon/brown paper bedding |
| | Compound discrimination - Extradimensional | 6 consecutive correct trials | Medium | Odor | Pebbles/paprika Pebbles/coconut | Sand/paprika Sand/coconut |

**Supplemental Table 2.** Correlational analysis between active errors during acquisition in R2 and pair-wise ROI connectivity (* P < 0.05).

| ROI pairs | R | P |
|---|---|---|
| CPu-HippD | -0.01 | 0.95 |
| CPu-IL | -0.24 | 0.13 |
| CPu-NAc | -0.39 | **0.01*** |
| CPu-OFC | -0.39 | **0.01*** |
| CPu-PrL | -0.19 | 0.23 |
| CPu-S1 | -0.07 | 0.68 |
| CPu-Thal | -0.12 | 0.47 |
| HippD-IL | -0.14 | 0.37 |
| HippD-NAc | -0.28 | 0.07 |
| HippD-OFC | -0.30 | 0.06 |
| HippD-PrL | -0.09 | 0.59 |
| HippD-S1 | 0.09 | 0.58 |
| HippD-Thal | 0.08 | 0.61 |
| IL-NAc | -0.16 | 0.33 |
| IL-OFC | -0.18 | 0.25 |
| IL-PrL | -0.16 | 0.31 |
| IL-S1 | -0.16 | 0.33 |
| IL-Thal | -0.27 | 0.09 |
| NAc-OFC | -0.21 | 0.19 |
| NAc-PrL | -0.25 | 0.11 |
| NAc-S1 | -0.25 | 0.11 |
| NAc-Thal | -0.36 | **0.01*** |
| OFC-PrL | -0.30 | 0.06 |
| OFC-S1 | -0.27 | 0.08 |
| OFC-Thal | -0.39 | **0.01*** |
| PrL-S1 | -0.09 | 0.56 |
| PrL-Thal | -0.29 | 0.06 |
| S1-Thal | -0.05 | 0.73 |

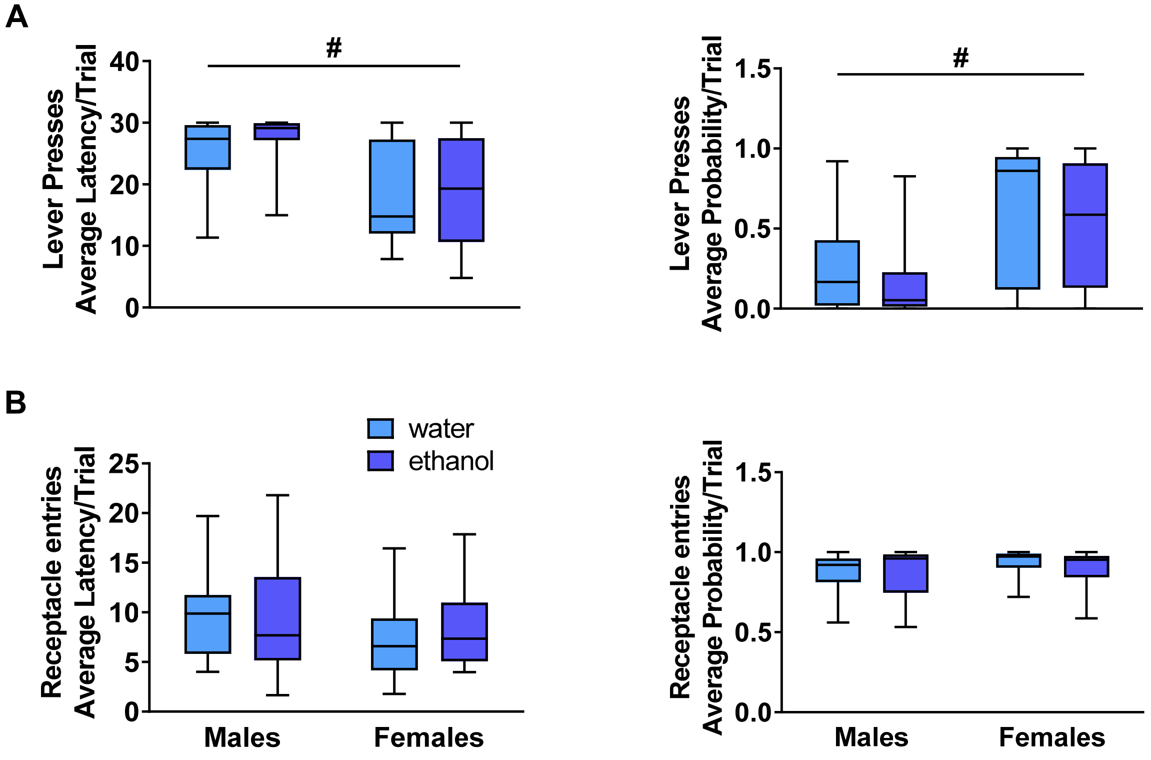


**Supplemental Figure 1. Sex influenced lever press metrics in Pavlovian conditioned approach. A.** Left: Female rats pressed the lever sooner than males (P<0.01). Right: Female rats also presented a higher probability of lever pressing than males (P<0.01). **B.** Groups did not significantly differ on receptacle latency (left) or probability of receptacle entry (right). Since the data were not normally distributed, data are presented in box plots showing median (horizontal line), interquartile range (box), and minimum and maximum data values (lower and upper whiskers). Detailed statistical analyses are provided in Table 1. # main effect of sex.

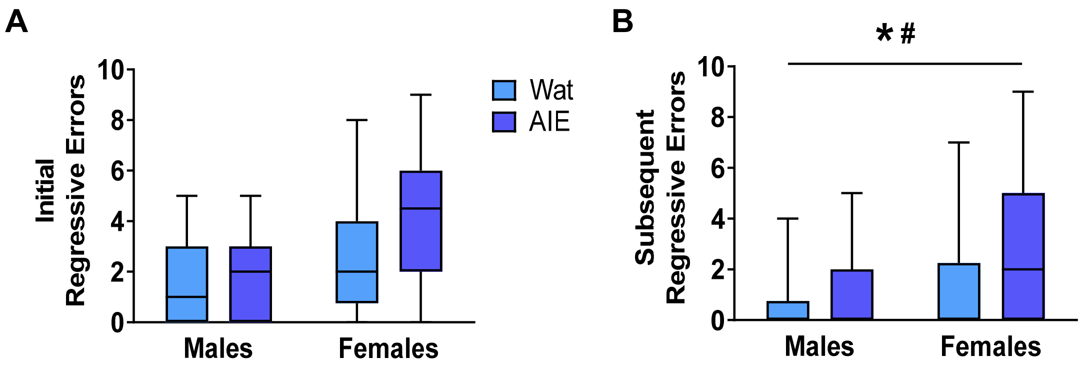


**Supplemental Figure 2. AIE exposure and sex influenced subsequent regressive errors in Reversal 2**. **A.** Groups did not differ in the number of initial regressive errors during acquisition of Reversal 2. **B.** Both AIE exposure (P=0.05) and female sex (P< 0.01) were associated with more subsequent regressive errors. Box plots show median, interquartile range and minimum and maximum data values. Detailed statistical analyses are provided in Table 1. # main effect of sex. * main effect of exposure.

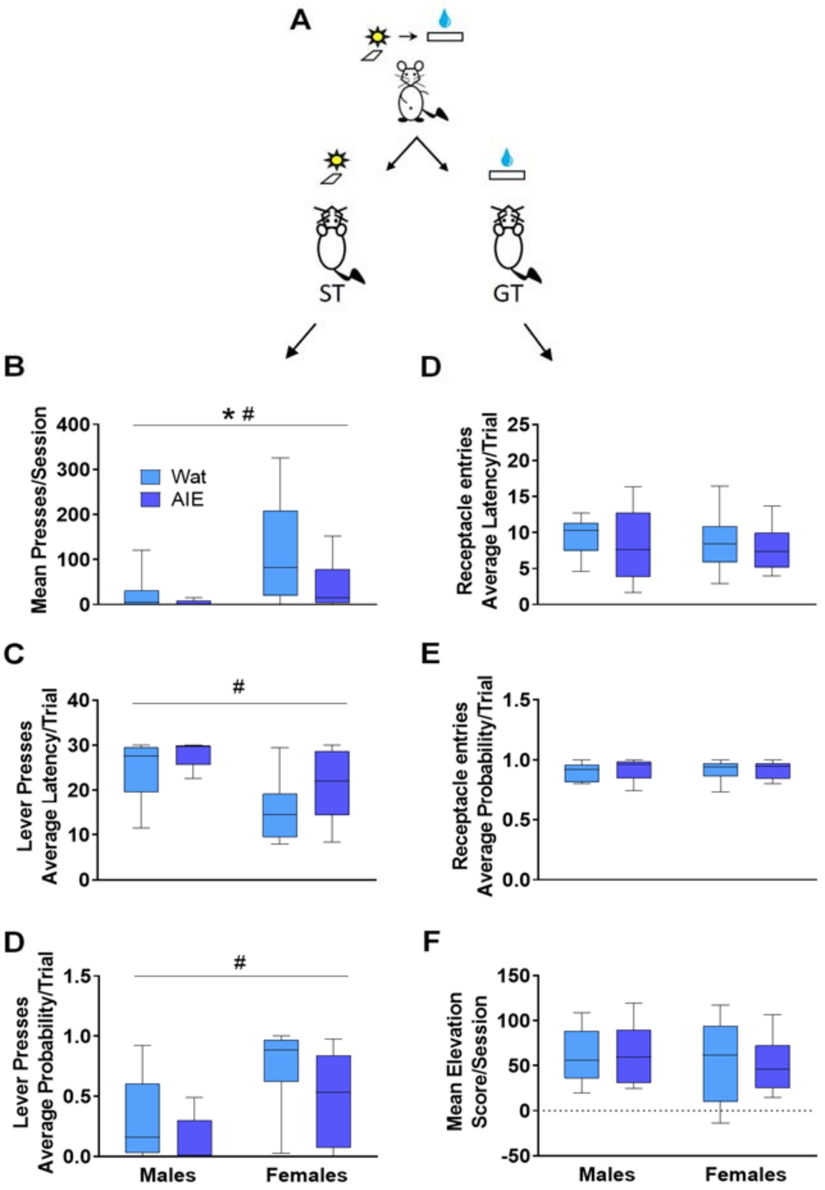


**Supplemental Figure 3. In the subset of rats undergoing fMRI, AIE exposure did not promote sign-tracking behavior. A.** Scheme representing phenotypes of interest in Pavlovian conditioned approach. During training, a compound-cue (light/lever) was presented during a 30-second period, followed by 100 µl of 20% sucrose that served as a reward. Sign-tracking (ST) rats preferentially interacted with the cue (light/lever) while goal-tracking (GT) animals preferentially interacted with the reward receptacle. **B.** Females rats pressed the lever more times than males (OR=10.630 [95% CI: 3.323 to 33.998], P<0.001). Interestingly, AIE-exposed animals pressed the lever fewer times than water controls (OR=5.862 [95% CI: 1.513 to 22.709], P=0.01). No interaction between exposure and sex (P=0.31) was observed. **C-D.** Female rats pressed the lever faster (OR=0.751 [95% CI: 0.593 to 0.950], P=0.02) and with a higher probability (OR=3.474 [95% CI: 1.316 to 9.167], P=0.02) than males. Neither metric yielded a significant main effect of exposure or sex-by-exposure interaction (P>0.05). **D-F.** No significant main effects or interactions were observed in any of the goal-tracking metrics (P>0.05). Since our data were not normally distributed, figures are presented using box plots and show median, interquartile range and minimum and maximum data values. # main effect of sex. * main effect of exposure. Female-AIE (n=10); female-water (n=12); male-AIE (n=9); male-water (n=10).

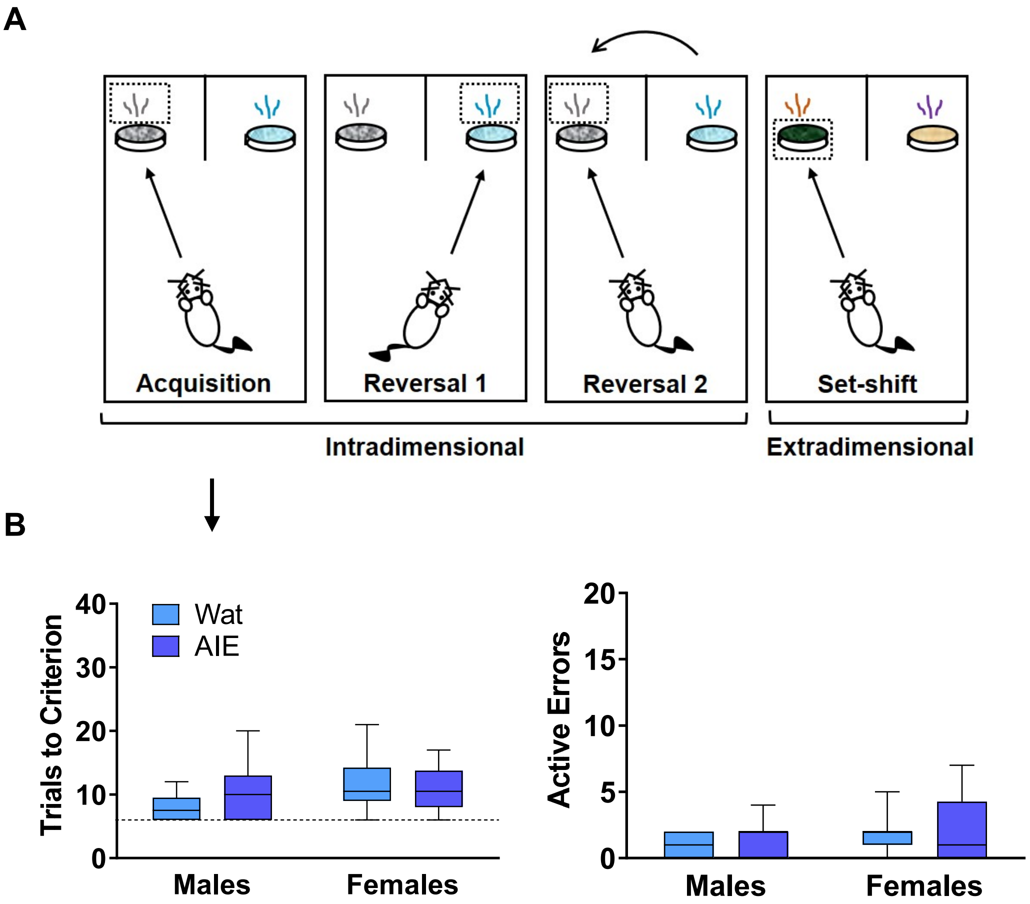


**Supplemental Figure 4. In the subset of rats undergoing fMRI, AIE exposure did not impair initial discriminative learning in a set-shifting task. A.**Scheme of the attentional set-shifting task. We trained rats to discriminate between two different odors, one of which was associated with a food reward (details about task in Supplemental Table 1). Cups containing rewards were covered with digging media, and animals were trained to dig inside the container according to the odor predicting the food, independent of the digging medium covering the cups. After initial training, intradimensional reversals 1 and 2 were introduced, keeping odor as the relevant dimension. Lastly, the extradimensional set-shift phase was initiated with novel stimuli, when the appropriate discriminant was the medium instead of odor. Criterion was set at 6 consecutive correct choices. **B.** Neither AIE exposure nor sex affected the total number of trials that animals required to reach criterion (6 correct consecutive choices; $P>0.05$) or the number of errors made ($P>0.05$). Moreover, no interaction was observed for any parameter ($P>0.05$). Female-AIE (n=10); female-water (n=12); male-AIE (n=9); male-water (n=10). Arrow indicates that graphs in panel B describe acquisition phase. Box plots represent median, interquartile range and minimum and maximum data values.

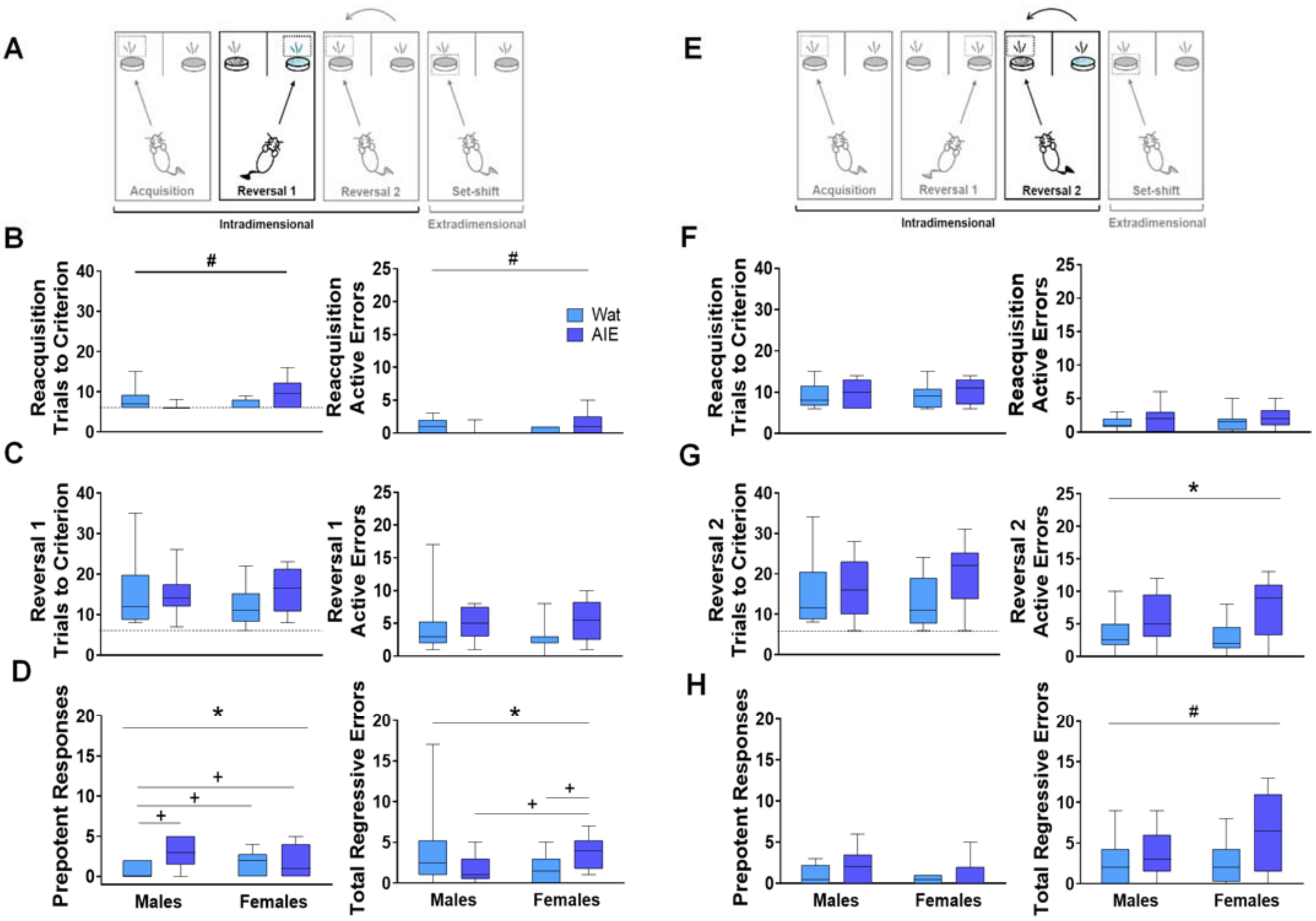


**Supplemental Figure 5. In the subset of rats undergoing fMRI, AIE exposure impaired some parameters in reversal learning. A.** On Reversal 1, the contingency was reversed such that the opposite odor signified the reward location. **B.** Every new training day began with reacquisition of the association from the previous day (reacquisition). Left: Female rats required more trials to reach criterion than males (OR=1.527 [95% CI: 1.097-2.124], P=0.01), with no main effect of exposure (P=0.11). We also identified a significant interaction between sex and exposure (OR=0.546 [95% CI: 0.348-0.856], P<0.01), but no pair-wise comparisons were significantly different after Bonferroni correction. Right: Similar to total trials, female rats made more errors in reacquisition (OR=7.2 [95% CI: 1.022-50.714], P=0.05), with no main effect of exposure (P=0.13). We observed a significant interaction between sex and exposure (OR=0.745 [95% CI: 0.007-0.49950], P=0.01), but no pair-wise comparisons were significantly different after Bonferroni correction. **C.** During Reversal 1, groups did not differ in the total number of trials required to reach criterion (left; Exposure: P=0.94; Sex: P=0.63; Exposure-by-Sex: P=0.24). Right: Similarly, no differences were observed in the number of active errors while reaching criterion (right; Exp: P=0.91; Sex P=0.50; or Exposure-by-Sex: P=0.26). **D.** As active errors involve different types of errors, we subdivided them into “prepotent” and “regressive” depending on whether they occurred before or after a correct response, respectively. Left: In general, AIE-exposed animals made more prepotent responses than water-exposed (OR=0.200 [95% CI: 0.072-0.556], P<0.01). For prepotent responses, we observed a significant interaction between exposure and sex (OR=4.167 [95% CI: 1.168-14.858], P=0.03). Post-hoc comparisons showed increased prepotent responses in AIE-exposed males when compared with male and female water controls (P<0.001 and P=0.04, respectively). Also, control females exhibited increased prepotent responses compared with control males (P=0.04). Right: We also observed an interaction between sex and exposure in regressive errors

(OR=0.210 [95% CI: 0.069-0.637], P<0.01). Post-hoc comparisons showed that AIE-exposed females made more regressive errors than control females (P=0.01) and AIE-exposed males (P=0.02). **E.** The next day, rats underwent reacquisition followed by Reversal 2, wherein the association was switched back to the original contingency used on Acquisition. **F.** In reacquisition of the previous day's rule, groups did not differ in the total number of trials required to reach criterion (left; Exposure: P=0.62; Sex: P=0.70; Exposure-by-Sex: P=0.66) or the number of active errors (Sex: P=0.34; Exposure: P=0.69; Exposure-by-Sex: P=0.98). **G.** When Reversal 2 was introduced, groups did not differ in the total number of trials required to reach criterion (left; Exposure: P=0.35; Sex: P=0.67; Exposure-by-Sex: P=0.27). AIE-exposed animals committed more total errors (right) than water-exposed rats (OR=0.611 [95% CI: 0.400-0.933], P= 0.02), but no significant main effect of sex (P=0.23) or exposure-by-sex interaction was observed (P=0.15). **H.** Regarding error type, prepotent responses were not different among groups (Exposure: P=0.12; Sex: P=0.19; Sex*Exposure P=0.94). The number of regressive errors was higher in female rats (OR=1.668 [95% CI: 1.099-3.531], P= 0.02), with no significant main effect of exposure (P=0.15) or exposure-by-sex interaction (P=0.09). Results in Reversal 1 and Reversal 2 suggest that AIE-exposed animals had more difficulty than water-exposed controls to inhibit learned rules, update learned information, and guide behavioral choices based on feedback. Those effects were also mainly observed in females rather than males. Box plots show median, interquartile range and minimum and maximum data values. # main effect of sex. * main effect of exposure. + simple differences after post-hoc pairwise comparisons. Female-AIE (n=10); female-water (n=12); male-AIE (n=9); male-water (n=10).

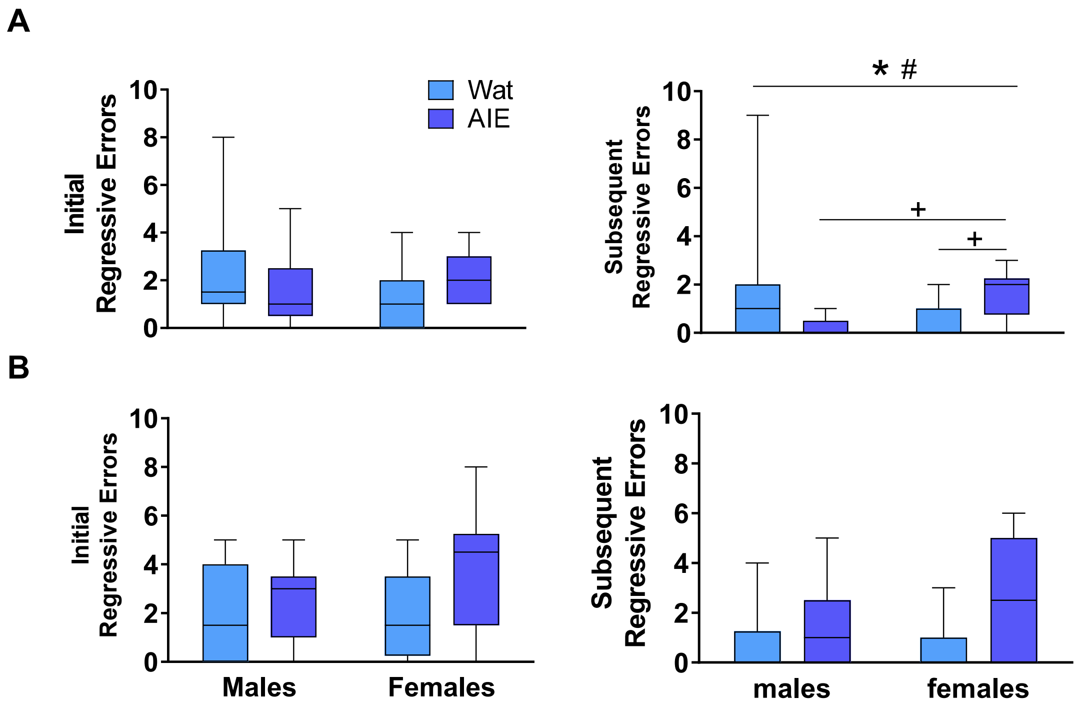


**Supplemental Figure 6. In the subset of rats undergoing fMRI, AIE exposure and sex influenced subsequent regressive errors in Reversal 1**. **A.** During Reversal 1, initial regressive errors were not impacted by AIE exposure (P=0.37) or Sex (P=0.40), and no significant exposure-by-sex interaction was observed (P=0.07). However, AIE-exposed rats made more subsequent regressive errors than did controls (OR=7.650 [95% CI: 1.130-51.807], P= 0.04) and females made more errors than did males (OR=7.2 [95% CI: 1.057-49.066], P= 0.04). We also observed an exposure-by-sex interaction (OR=0.048 [95% CI: 0.005-0.446], P< 0.01). Bonferroni-corrected post-hoc comparisons confirmed that female AIE-exposed animals committed more subsequent regressive errors than female controls (P<0.01) and AIE-exposed males (P<0.001). **B.** During Reversal 2, groups did not differ on the number of initial regressive errors or subsequent regressive errors. Box plots show median, interquartile range and minimum and maximum data values. # main effect of sex. * main effect of exposure. + simple differences after post-hoc pairwise comparisons. Female-AIE (n=10); female-water (n=12); male-AIE (n=9); male-water (n=10).

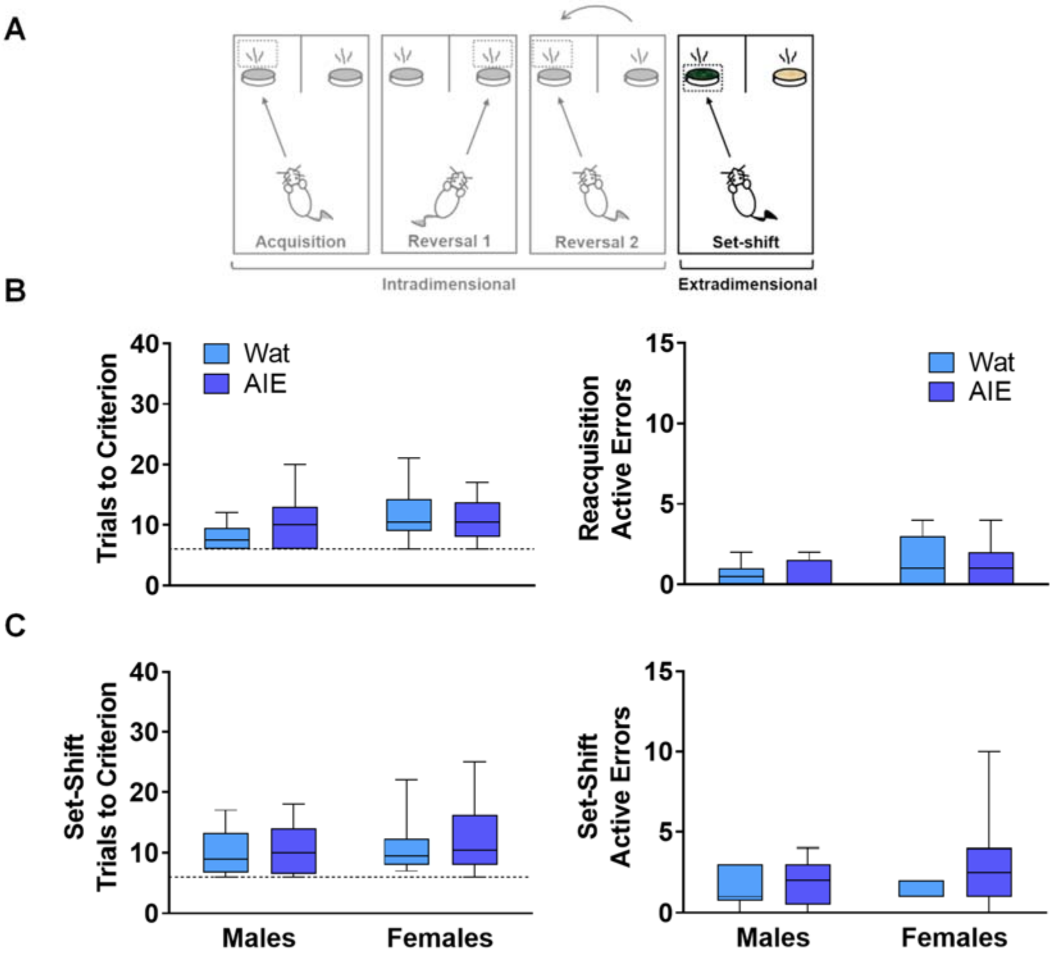


**Supplemental Figure 7. In the subset of rats undergoing fMRI, AIE exposure did not impair set-shifting performance. A.** After reversals, an extradimensional set shift to completely new stimuli and sensory modality of the discriminant was introduced. In this phase, the discriminant associated with the reward was now the digging medium and odor was irrelevant. **B.** In the reacquisition phase, groups did not differ in the total trials to reach criterion (left; Exposure: P=0.85; Sex: P=0.42; Exposure*Sex: P= 0.92) or active errors (right; Exposure: P=0.85; Sex: P=0.32; Exposure*Sex: P= 0.60). **C.** In the extradimensional set shift, groups did not differ in the total trials required to reach criterion (left; Exposure: P=0.60; Sex: P=0.26; Exposure*Sex: P= 0.69) or the number of active errors made (right; Exposure: P=0.51; Sex: P=0.17; Exposure*Sex: P= 0.20), suggesting that AIE exposure does alter the ability to shift sensory modality of the discriminant in the context of new stimuli. Box plots show median, interquartile range and minimum and maximum data values. Female-AIE (n=10); female-water (n=12); male-AIE (n=9); male-water (n=10).
